# Population structure amplifies mutation load

**DOI:** 10.1101/2025.09.04.674237

**Authors:** Nikhil Sharma, Suman G. Das, Arne Traulsen, Joachim Krug

**Affiliations:** Department of Theoretical Biology, Max Planck Institute for Evolutionary Biology, August-Thienemann-Str. 2, 24306 Plön, Germany; Institute for Ecology und Evolution, University of Bern, 3012 Bern, Switzerland; Swiss Institute of Bioinformatics, 1015 Lausanne, Switzerland; Institute for Biological Physics, University of Cologne, D-50937 Cologne, Germany

## Abstract

The interplay of mutation and natural selection is key to evolution: Mutations introduce genetic variation, whereas selection favours higher-fitness individuals. The balance between the two is quantified by the mutation load—the reduction in mean population fitness due to deleterious mutations. In large well-mixed populations, selection dominates, and the mutation load is proportional to the mutation probability per reproduction event. However, the role of spatial structure in shaping mutation–selection dynamics is poorly understood. Here, we use evolutionary graph theory to show that heterogeneous spatial structures generically amplify the effect of mutation relative to selection. In contrast to well-mixed populations, the mutation load in heterogeneous structures can remain substantial even when the mutation rate is very small, signaling a breakdown of selection. For example, in star-shaped populations the mutation rate is effectively amplified by a factor proportional to the population size. When mutation is coupled to reproduction, this leads to the proliferation of less-fit types over fitter ones, a scenario of “survival of the weakest”. The reduction of natural selection may help explain, for example, tumour heterogeneity in spatially structured tissues. It also suggests that heterogeneous social networks can promote the persistence of unpopular opinions, thereby fostering diversity. By showing how spatial structure modulates the effect of selection, our work points to potentially new ways of steering evolution toward desired outcomes.

## Introduction

“Survival of the fittest” is among the most frequently used phrases in evolutionary biology. With only two competing individual types, survival of the fittest implies that the type with higher fitness eventually takes over the population due to natural selection. In the presence of mutations, a non-zero proportion of less fit individuals coexists with the fitter individuals—referred to as mutation-selection balance. The reduction of the population fitness induced by the presence of less-fit individuals is called the mutation load [1–3]. While the balance between natural selection and mutation has been extensively studied in well-mixed populations [4–6], broadly applicable results concerning this interplay in structured populations are rare. Yet, real-world populations are not well-mixed and often exhibit spatial structure—for example in cancer evolution [7–9] or the evolution of antibiotic resistance in microbes [10–12]. Recent studies have hinted that spatial structure may elevate mutation load [13, 14], but both the extent of this effect and its underlying mechanisms remain poorly understood. Here, we argue that population structure generally increases mutation load, and that this effect can be especially pronounced in strongly inhomogeneous structures. Crucially, we provide a mechanistic explanation for this phenomenon, demonstrating how structural asymmetries systematically distort the balance between mutation and selection.

We employ birth-death dynamics on graphs, a versatile and well-established framework for studying evolutionary dynamics in structured populations [15–17]. The population structure is modelled by a graph, with nodes representing individuals and links defining their neighborhood. During a reproduction step, a node can only be replaced by the offspring of a node in its neighborhood. This way, a well-mixed population is equivalent to the complete graph where any node can be replaced by any other node, making all nodes indistinguishable. In contrast, on a regular graph such as a two-dimensional lattice, each node has a fixed number of neighbors (four in the case of the square lattice) and can only be replaced by one of them. Figure 1 shows exemplary results obtained from simulations of mutation-selection dynamics on different graph structures. The panels display the frequency of the mutant type at mutation-selection balance as a function of its fitness *f* relative to the wild type. Natural selection increases the mutant frequency when it is beneficial (*f >* 1) and decreases it when the mutant is deleterious (*f <* 1). For all structures shown in Figure 1, natural selection is weakened, and hence the mutation load is amplified compared to the well-mixed case. The effect is moderate for the regular lattice and the random Erdős-Rényi graphs, and becomes more pronounced for the strongly heterogeneous Barabási-Albert graphs and Cayley trees.

**FIG. 1.**
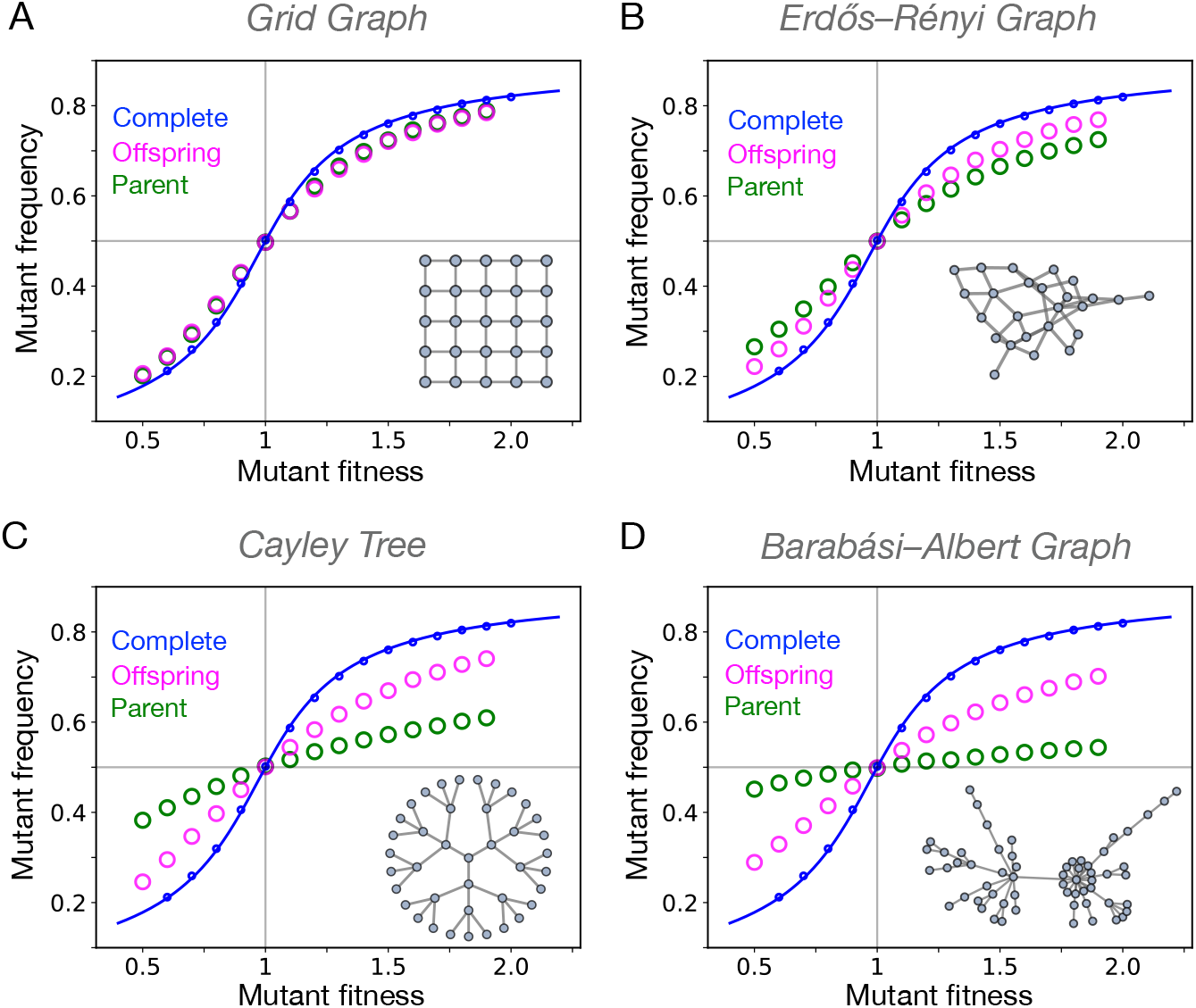
Population structure increases the mutation load. Steady-state mutant frequencies for the simulated mutation-selection process are shown for four graph families: a two-dimensional lattice with periodic boundary conditions (panel A), Erdős-Rényi graphs (panel B), a Cayley-tree with branching number 3 (panel C), and Barabási-Albert graphs generated by linear preferential attachment (panel D). In all cases the population consists of *N* = 100 individuals and mutations between the two types occur symmetrically with probability *µ* = 0.1 per reproduction event. Compared to the well-mixed population (blue curve), all graph families exhibit a higher mutation load. Beneficial mutants are less abundant in structured populations than in the well-mixed population due to weakened natural selection, and deleterious mutations are more abundant. Furthermore, the parent movement rule (green markers) results in a higher mutation load than the offspring movement rule (magenta markers). The increase in mutation load is most pronounced for the strongly degree-inhomogeneous Barabási-Albert graph and the Cayley tree equipped with parent moving dynamics. A reversal in the mutation-selection profile—where deleterious mutants outnumber beneficial ones—is observed in Barabási-Albert graphs generated via nonlinear preferential attachment (see App. A3).

Additionally, the mutation load is seen to depend substantially on the update rule used in the simulations. Previous work on evolutionary dynamics on graphs has shown that the results are often surprisingly sensitive to model details [18–21], which makes it challenging to arrive at general statements. For example, one time step of the commonly used Moran dynamics consists of a birth event (where an individual is chosen for reproduction) and a death event (where a neighboring individual is removed in order to keep the population size constant). Depending on the order in which the two steps are applied one distinguishes between Birth-death (Bd) and death-Birth (dB) dynamics, where the upper case letter marks the event (here birth) where selection acts and the lower case letter (here death) represents a neutral event. Studies investigating the fixation dynamics following the introduction of a single mutant have shown that most population structures amplify natural selection over genetic drift under Bd updating, but suppress selection for dB updating [22].

Recently it has been pointed out that not only the order of birth and death events, but also the type of individual replacing the dead individual can strongly affect the dynamics, because it determines the local environment of the mutant individuals and hence their evolutionary fate [23]. This is important in the present context, because any model describing mutation-selection dynamics in structured populations needs to specify the placement rules for the mutated offspring. It is a frequent assumption in the literature that the mutant offspring moves, but there is no a priori reason why movement of one kind shall be preferred over another. For example, in symmetrically dividing unicellular populations, non-mutant offspring are as likely to move as mutated ones, provided the mutation does not affect mobility. This is of relevance to clinical applications such as the modelling of cancer cell dynamics in tumors. Due to strong population regulation in tumors, death and birth events occur at the same rates justifying using birth-death models [8, 24–26], but the frequent assumption that the mutant offspring moves [27] can lead to biased predictions of mutation load. Similar concerns apply to microbial evolution in structured environments. Nevertheless, most mathematical models assume the movement of the mutant daughter cell [25, 26]. We call this offspring movement and contrast it with the parent moving rule, where the parent-resembling daughter cell moves and replaces the dead individual. Figure 1 shows that parent movement can strongly enhance the mutation load in structured populations. In fact, for the Barabási-Albert graphs, the effect of natural selection is almost obliterated, as the mutant frequency is pushed close to the neutral limit of ^1^.

We systematically investigate the mutation-selection balance in graph-structured populations, addressing specifically the role of parent moving dynamics vs. offspring moving dynamics. We also ask to what extent the results are robust to different birth-death update schemes. To tackle these questions, we study birth-death processes with mutation on different population structures, and compare the respective mutation-selection balance state with that of the well-mixed population. We show that allowing parent individuals to move in place of the offspring can dramatically increase the mutation load across different families of heterogeneous graphs. In some cases, the mutation load becomes independent of the mutation probability *µ* and remains nonzero in the limit *µ*→0, a striking phenomenon that we designate as strong mutation load amplification. As a paradigm of a strongly inhomogeneous graph we use the star graph, which allows for thorough analysis and detailed understanding of the dynamics on a highly heterogeneous population structure. Additionally, we show the robustness of our results under death-birth update schemes and scenarios where mutations arise spontaneously in the population rather than being coupled to birth events.

## Results

Consider a population of two types of individuals, mutant-type and wild-type, where the mutant-type has fitness *f* relative to the wild-type. In a given update step, an individual is selected with a probability proportional to its fitness to reproduce. The offspring can mutate to the other type with probability *µ*. After that, a neighboring individual of the parent is randomly chosen for death. Then, either the offspring replaces the dead individual or the parent individual moves, with the offspring occupying the parent node. We use the shorthand Bd^*o*^ for the scenario where offspring moves, and Bd^*p*^ for the parent moving case. Note that this choice matters only if a mutation has occurred.

The mutant frequency in a well-mixed population is obtained by solving the discrete time replication-mutation dynamics, see App. A1 for details. Assuming that the mutant is the fitter type (*f >* 1), the maximal possible population fitness is *f*, and correspondingly the mutation load in steady state is [28]

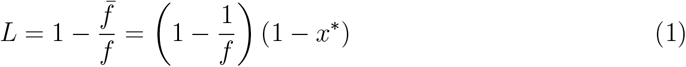

where *x*^*\**^ is the mutant frequency in steady state and 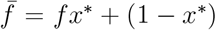 the steady state population fitness. The mutation-selection balance on the complete graph (or, equivalently, a well-mixed population) is identical for the two update rules Bd^*o*^ and Bd^*p*^. In the limit of small mutation rate, *µ* → 0, the mutation load of a well-mixed population is *L* = *L*_*C*_ ≈ *µ* independent of the selection strength, a classic result due to Haldane [1, 4].

The simulation results for four canonical graph families presented in Figure 1 show that these four population structures have a higher mutational load (higher frequency of the deleterious type) than the well-mixed population. Specifically, parent movement generates higher accumulation of deleterious individuals than the offspring moving dynamics. Furthermore, greater variation in the graph’s degree distribution increases the difference between the steady-states of the parent moving and offspring moving dynamics.

For a first qualitative understanding of these effects, let us consider Bd dynamics, where an individual is chosen for reproduction with probability proportional to its fitness. In a graph with varied degree distribution, highly connected nodes get replaced more often and hence have a higher turnover rate. In the absence of mutations, fitter individuals—which are more likely to be chosen for reproduction—preferentially reproduce into high-degree nodes. When mutations occur, the offspring moving dynamics will tend to place the mutated offspring in high-degree nodes, but the high turnover will ensure that deleterious mutants do not persist for long in these nodes. In contrast, the parent moving dynamics will tend to move the selected fitter parents to high degree nodes, and leave behind less-fit mutant offspring in lower degree nodes, where they will persist longer due to low turnover rates. In short, for parent moving dynamics, the occupancy of high-degree nodes is predominantly changed by positive selection and that of low degree nodes is predominantly changed through the placement of deleterious mutations. In this scenario, a higher abundance of low degree nodes can lead to a higher mutational load in the population. In the Cayley tree in Figure 1C, nearly 2/3 of the nodes in population are peripheral nodes, which have a lower degree than the internal nodes, leading to a substantial increase in the load under the parent-moving rule. For the Barabási-Albert graph, the heavy tail degree distribution implies that high degree ‘hubs’ are rare and most nodes have relatively low degree, enabling the parent moving dynamics to cause an even stronger amplification of mutation.

To validate these arguments, we investigate the offspring and parent Birth-death dynamics on the star graph where all leaf nodes are connected to one central node. Under offspring moving dynamics (Bd^*o*^), the central node of the star graph changes its state once a leaf node is chosen for reproduction and the leaves change their state once the central node is chosen for birth. Thus, the central node changes its state *N* times faster than the leaves, and for large *N* it effectively reaches a quasi steady-state before the leaves can change. A detailed analysis shows that this time-scale separation generates the same mutation-selection balance state for the center and the leaf nodes as the complete graph, see Figure 2A and App. A2a. Therefore, the star graph under Bd^*o*^ dynamics has the same mutation load as the well-mixed population. This result was noted earlier [29], but it is an exception rather than the rule: Under all other updates, the star graph is a mutation load amplifier.

**FIG. 2.**
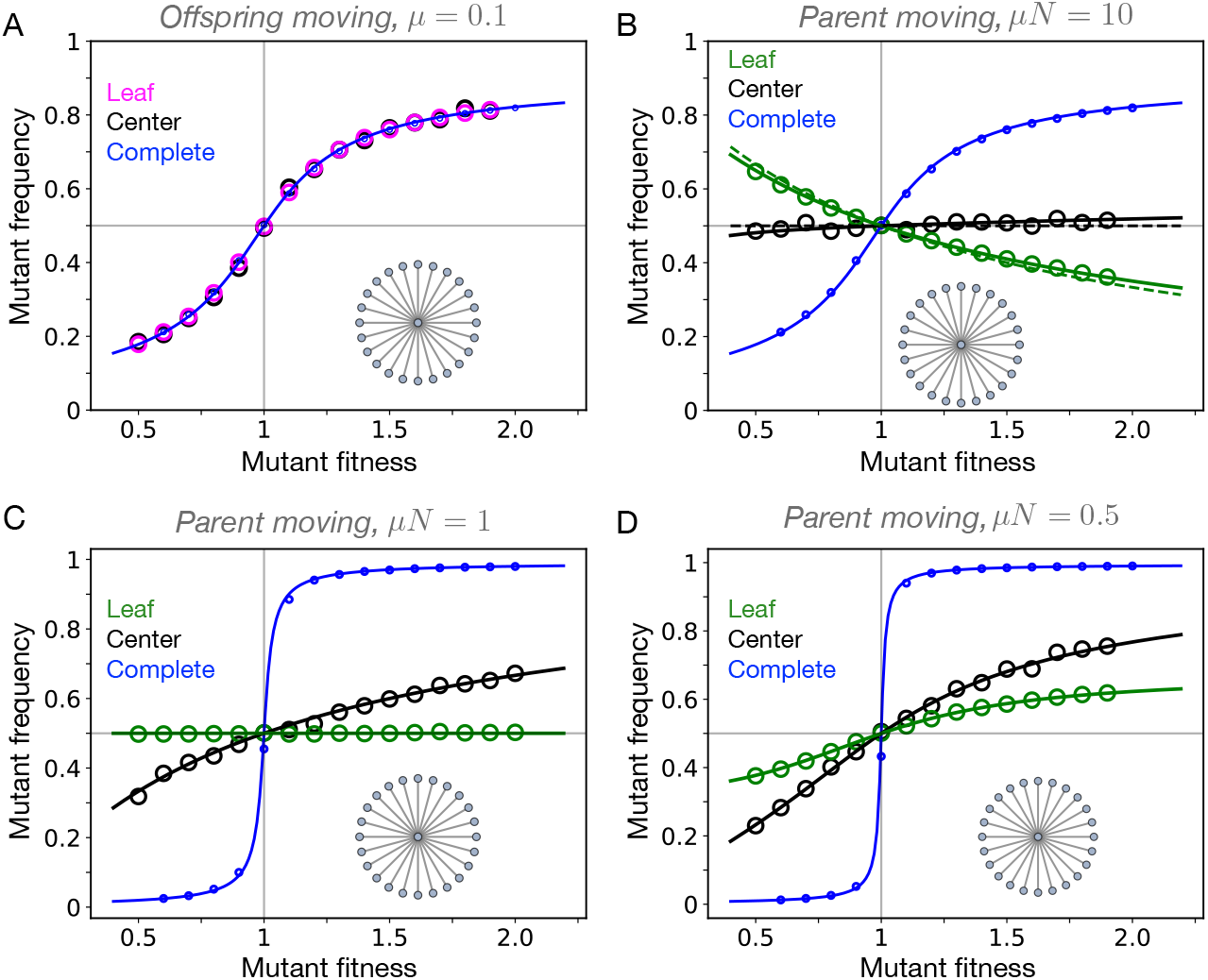
Birth-death dynamics on the star graph. A: Under the Bd update rule with offspring movement (Bd^*o*^), the star graph exhibits the same mutation-selection balance as the complete graph. Both the mutant frequency at the leaf nodes (magenta markers) and at the central node (black markers) are identical to those observed at mutation-selection balance of the complete graph (small blue circles and full line). The mutation probability is *µ* = 0.1 and the population size *N* = 100. B-D: In contrast to the offspring-moving case, the mutation-selection balance for the star graph under the Bd parent-moving update rule (Bd^*p*^) depends on the product *µN*. The three panels show results for a population of size *N* = 100 and mutation probabilities *µ* = 0.1 (B), *µ* = 0.01 (C) and *µ* = 0.005 (D). The mutant frequency for the complete graph is shown in blue. In panel B, deleterious mutants are found at a higher frequency than beneficial mutants: Parent moving dynamics inverts natural selection. The dashed green line shows the large *N* approximation given by Eq. (3), while the solid lines represent the exact analytical solution (see App. A2b). In panel C, the steady-state leaf mutant frequency in the star graph is independent of the mutant’s fitness. Here selection is perfectly counter-balanced by mutations. In panel D, natural selection is revived and beneficial mutants have higher frequencies than the deleterious ones. Overall, the mutational force dominates natural selection in the star graph under Bd^*p*^ updating.

For the case when the parent moves, a leaf node can change its state with probability *µ* every time it is chosen for reproduction. This is contrary to the case of offspring moving dynamics, where leaf nodes can only be changed if the center is selected for reproduction. Thus, for large *N* the changes in leaves are mostly driven by mutations. The leaf dynamics gets effectively decoupled and becomes independent of the center state. We can therefore treat each leaf node as an isolated node experiencing Bd^*p*^ dynamics. Let 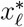 be the probability that a leaf node is occupied by a mutant in the steady-state, and denote the probabilities that the node transitions from wild-type to mutant and vice versa by *T*_*w→m*_ and *T*_*m→w*_, respectively. In the stationary state 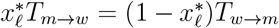 and therefore

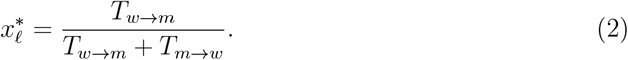

As nodes are selected for reproduction according to fitness, *T*_*m→w*_*/T*_*w→m*_ = *f*, which yields

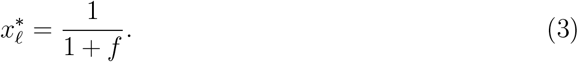

The lower the mutant’s fitness, the more likely it is that a leaf node is occupied by a mutant. In other words, mutation not only reduces natural selection but actually reverts its direction. Since all the leaf nodes are identical and almost all nodes are leaves for large *N*, 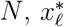 is the steady-state mutant frequency in the population. In Figure 2 B, we see that (3) provides a good approximation to the average mutant leaf frequency.

Using Eqs. (1) and (3), the mutation load on the star graph under parent moving dynamics turns out to be 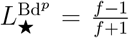. Unlike the well-mixed population, the mutation load is independent of *µ* and remains nonzero even in the limit *µ* → 0. The detailed analysis in App. A2b shows that Eq. (3) is exact for the star graph with parent movement for any *µ >* 0 and *N* → ∞. Note that a reversal of natural selection due to mutations, as seen in Figure 2B, occurs also in the well-mixed case, but only if the mutation probability per reproduction step exceeds 1/2 (see App. A). It is ultimately linked to the coupling of mutations to reproduction, which causes fit individuals to produce mutated offspring more frequently than unfit ones. In the star graph, the effect of mutations is massively amplified because the equations used to obtain steady-state frequencies contain the product *µN* instead of *µ*. The appearance of the population-wide mutation rate *µN* in the condition for mutation-selection balance is the mathematical hallmark of the strong amplification of mutation load in the star graph. Natural selection is recovered only when the large population limit *N* → ∞ is taken simultaneously with a low mutation limit *µ* → 0 at fixed *µN*, see Figures 2 C,D.

How robust is the property of the star graph to strongly amplify the mutation load? To answer this question, we first conducted simulations where the offspring moving dynamics was adulterated with a small proportion of parent moving dynamics. We found that a proportion of 10% of parent moving update steps is sufficient to invert the mutation-selection profile, see Figure S3. Second, we investigated the mutation-selection balance on the star graph under death-Birth dynamics. For parent moving dynamics (dB^*p*^) the mutation load is again found to be strongly amplified with a steady-state mutant frequency 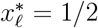 and a mutation load of 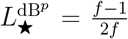 for large *N* (see App. A2d for details). For dB^*o*^ dynamics a full analytic solution for the steady state is not available because of a lack of separation of time scales (as for Bd^*o*^) or decoupling between mutation and selection dynamics (as for Bd^*p*^). Nevertheless a lower bound on the mutation load can be derived which shows that it is at least 2*µ* for small *µ*, and thus exceeds Haldane’s result for the well-mixed case.

At last, we relaxed the requirement for mutations to appear during reproduction events and considered a dynamics where mutations appear spontaneously with probability *µ* at a randomly chosen node. With probability 1 −*µ*, a Moran Bd or dB update without mutation takes place. We find that the star graph under Bd dynamics with spontaneous mutation is a strong amplifier of mutation load with *µ* replaced by *µN* and 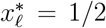 for large *N* (see App. B2a for details). For dB update, the spontaneous mutation dynamics is similar to the dB^*p*^ model, and the mutation load is at least 2*µ* for large *N* and small *µ*. Therefore, the amplification of mutation load by the star graph is robust to the proportion of offspring movement, the update rule and the mode of mutation. In Figure 3, we summarise the results for the mutation load on the star graph.

**FIG. 3.**
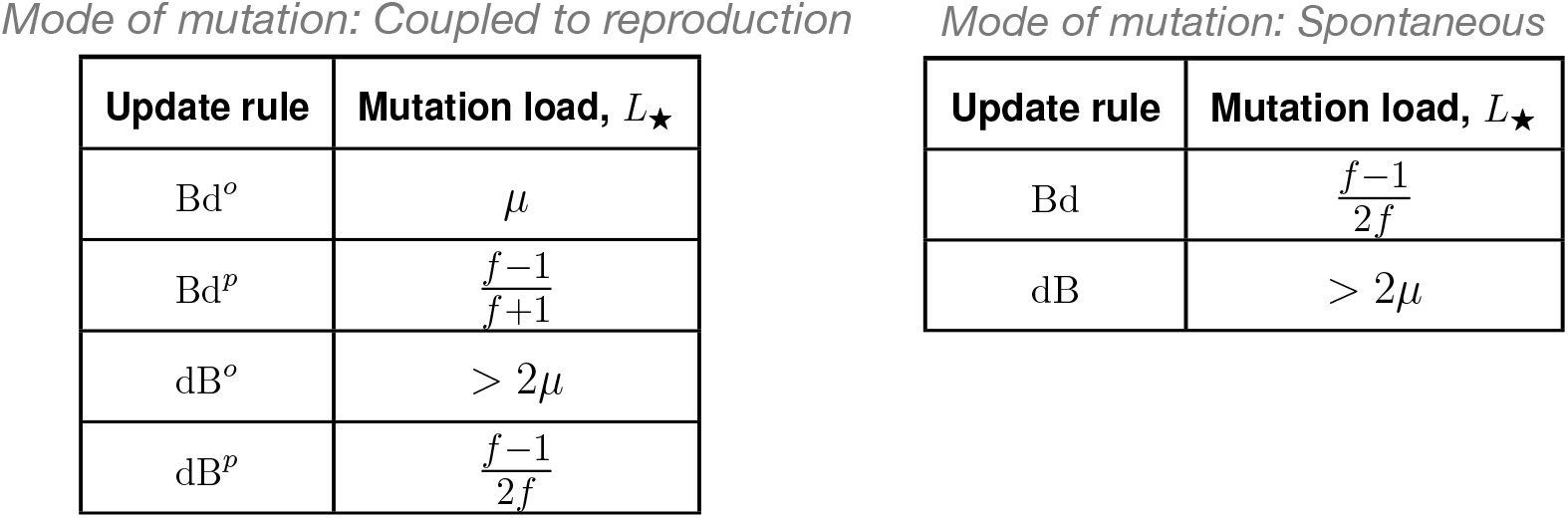
Mutation load on the star graph. The tables summarize the results for the mutation load on the star graph for different update schemes. In all cases, the leading order behavior of *L*_*_ for large *N* and small *µ* is presented. With the mutation load *L*_*C*_ = *µ* for the complete graph as a reference, we find that the mutation load on the star graph is higher than *L*_*C*_ for all cases except for Bd^*o*^. Thus, the star graph is an amplifier of mutation load. For the parent movement rules and for Bd update with spontaneous mutations, *L*_*_ becomes independent of *µ* for *µ* → 0, making the star graph a strong amplifier of mutation load.

## Discussion

By studying the interplay of mutation and natural selection in structured populations, we have found that heterogeneous population structures amplify mutation over natural selection. The frequency of disadvantageous individuals in structured population is higher than in the well-mixed population. The effect of mutation gets strongly enhanced when instead of the standard case of offspring moving, the parent individual moves after reproduction. In highly heterogenous graphs like the BA graph and the star graph, there is a reversal in the mutation-selection balance profile where deleterious mutants are found in higher abundance than the beneficial ones. On these graphs, we find “survival of the weakest” – contrary to survival of the fittest in a well-mixed population.

Weakened natural selection in heterogeneous spatial structures has broad implications. One key consequence is the ability of populations to evolve in rugged fitness landscapes [30, 31]. Populations with heterogeneous structures should be better at crossing fitness valleys than well-mixed populations due to their higher tolerance for deleterious mutants. Intratumour heterogeneity is another area where our results are relevant. The diversity of mutant strains in tumors is typically attributed to either nearly neutral mutations [32, 33] or ecological interactions among subclones [34, 35]. Our work adds a new perspective to this list by showing that heterogeneous spatial structures can contribute to tumor diversity by effectively neutralizing fitness differences among mutants.

Important applications of our findings can also be found in the social sciences, particularly in understanding the dynamics of opinions in social networks [36, 37]. In this context, the Bd model corresponds to the biased invasion process [38], where each node represents an individual holding one of several possible opinions. Popular opinions have higher “fitness” as they are more likely to be adopted by others. Previous studies have shown that highly heterogeneous structures favour the spread of popular opinions [39] — provided individuals do not change their opinions independently, such as through external influences or introspection. However, when individuals are allowed to change their opinions spontaneously, our findings indicate that heterogeneous social structures can also promote the persistence of less fit opinions, thereby increasing opinion diversity within the population. A recent theoretical study [40] focused on heterogeneous structures under neutral dynamics (and no mutation) also shows that heterogenous population structure can promote diversity, complementing our results.

What is next? An obvious open question is to quantify the mutation load amplification factor for heterogeneous networks such as the BA graphs in terms of the degree distribution. More fundamentally, our results indicate that all structures either amplify mutation load or have the same load as a well-mixed population. This suggests that the effect we describe is more robust than previously known effects of structure on evolutionary dynamics, for which both amplifiers and suppressors (of selection, fixation or diversity) have been found in all update categories [15, 19, 22, 23, 40–43]. We thus pose a question: Are there spatial structures that (contrary to the findings of this article) protect a population from the deleterious effects of mutations? This work provides a suitable platform where such research directions can be pursued.

## METHODS

The mathematical derivations of the mutation load for the complete graph (well-mixed population) and the star graph under various update schemes are presented in the supplementary information file.

## DATA AVAILABILITY

The data to reproduce our figures is available on Zenodo https://doi.org/10.5281/zenodo.15773813.

## CODE AVAILABILITY

The source code (Mathematica files / Jupyter notebooks) to reproduce our figures is available on Zenodo, https://doi.org/10.5281/zenodo.15773813.

## ACKNOWLEDGMENTS

We thank Claudia Bank, Michael Stumpf and Akanksha Singh for helpful discussions. We thank Carsten Fortmann-Grote for computational support. This work was supported by the Max Planck Society and by DFG within CRC 1310 “Predictability in Evolution”.

## AUTHOR CONTRIBUTIONS

N.S, S.G.D, A.T and J.K designed research; performed research; analyzed data; and wrote the paper.

## Supplementary Information

## Appendix A: Mode of mutation: coupled to birth

For mutations coupled to birth, mutations occur during reproduction events. We consider two types of individuals: wild-type and mutant-type. A mutant-type individual has a fitness *f* relative to a wild-type individual. We work with four Moran birth-death update rules: Bd^*o*^, Bd^*p*^, dB^*o*^, and dB^*p*^. The letters ‘b’ and ‘d’ represent the types of events—’b’ for birth and ‘d’ for death. If selection acts during an event, it is denoted by an uppercase letter. The notation also indicates the order for events. Finally, the superscript denotes the type of individual moving to vacant sites—’p’ for parent-type and ‘o’ for offspring-type.

For illustration, let us consider the example of the Bd^*o*^ update rule (see Fig. S1):

- Birth (B): An individual is selected to reproduce with a probability proportional to its fitness.
- death (d): A neighbouring individual of the reproducing individual is chosen at random with uniform probability to die.
- Replacement by offspring (*o*): Finally, the offspring replaces the dead individual, completing the update step.

The offspring either resembles the parent-type with probability 1 −*µ*, or mutates to another type with probability *µ*. Repeating these steps a sufficiently large number of times results in a steady-state for the Bd^*o*^ dynamics with mutations. The same process applies to other update schemes.

### 1. Well-mixed population: Complete graph

In the absence of mutations, advantageous mutants (*f >* 1) introduced into a very large well-mixed population of wild-types (*f* = 1) will eventually take over the population. This phenomenon is commonly referred to as *survival of the fittest*. However, when mutations are present, the long-term state of the population includes a finite frequency of wild-type individuals, and the population exhibits a *mutational load*. Mutational load results from the balance between natural selection, which promotes the spread of the fittest type, and mutation, which introduces variation.

Let us consider Bd^*o*^ updating to study mutation-selection dynamics on a complete graph. Denoting *x* as the frequency of the mutant-type and *y* as the frequency of the wild-type, we have *x* + *y* = 1. Therefore, we only need to track the frequency of one type. The expected change in mutant frequency after one Bd^*o*^ update step is,

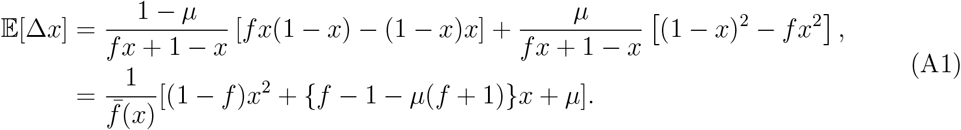

Where 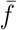 is the mean population fitness. In the steady-state, 𝔼 [Δ*x*] = 0. Using Eq. A1, the solution for the steady-state mutant frequency for *f ≠* 1 is,

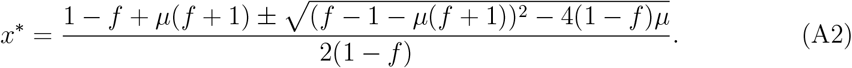

We focus on the second root, as the first root gives *x*^*\**^ outside the [0, 1] interval. For small *µ*, the solution can be approximated as,

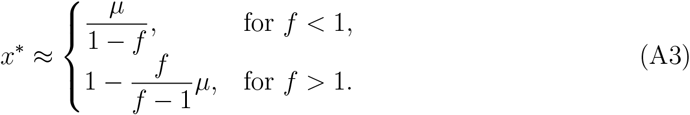

At neutrality (*f* = 1), Eq. A1 gives *x*^*\**^ = 1*/*2, as expected. For low *µ*, beneficial mutants are abundant compared to deleterious mutants, as shown in Fig. S2 A. Assuming advantageous mutant-type, Eq. A3 gives the mutational load for the well-mixed population,

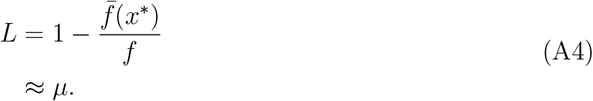

This is the well-known Haldane’s principle.

As *µ* increases, the proportion of deleterious mutants in the population is expected to rise, while the steady-state mutant frequency of beneficial mutants decreases. Thus, there must be an intermediate mutation probability where natural selection is precisely countered by mutation, ensuring *x*^*\**^ = 1*/*2 regardless of mutant fitness. Mathematically, this corresponds to a value of *µ* where 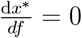. As expected, at *µ* = 1*/*2, the derivative of *x*^*\**^ with respect to mutant fitness *f* vanishes. Expanding the exact solution Eq. A2 near *µ* = 1*/*2, we obtain

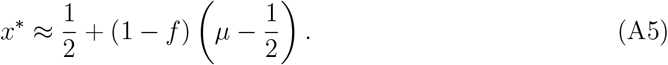

For *µ >* 1*/*2, the second term in Eq. A5 is positive for deleterious mutants and negative for beneficial mutants. As a results, deleterious mutants exceed the wild-type in abundance, whereas beneficial mutants become less frequent, as shown in Fig. S2 B. Even in the absence of an initial mutation bias, coupling mutations with birth introduces an implicit bias at higher mutation probabilities, leading to a reversal in the mutation-selection balance profile. For large *µ* (*µ* ≈ 1), the mutant frequency *x*^*\**^ at mutation-selection balance is given by

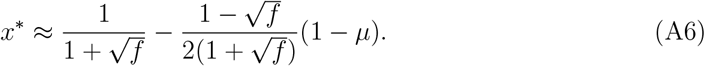

**Fig. S1.**
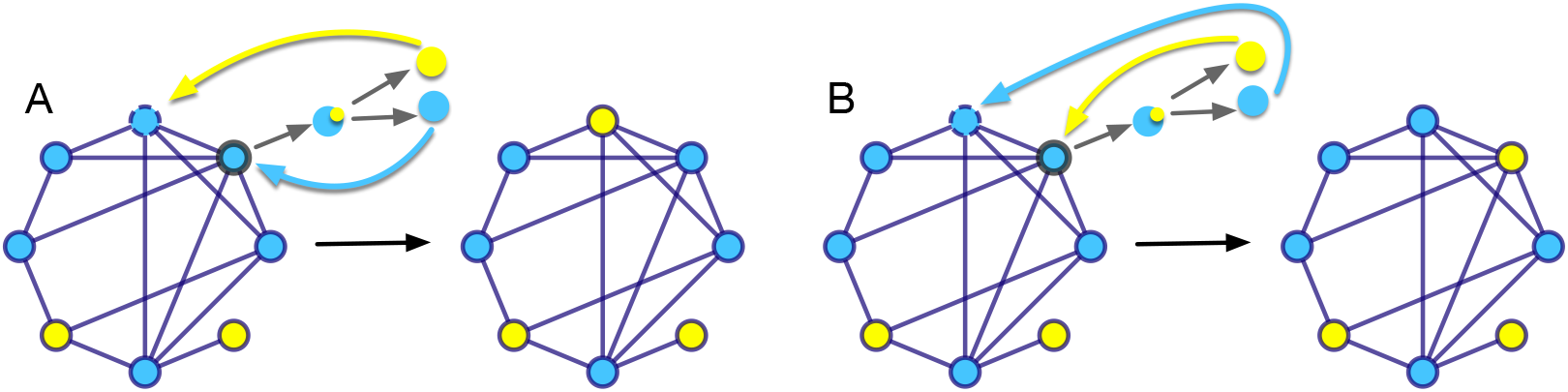
Moving offspring or parent? The population consists of wild-type (blue) individuals and the mutants (yellow), and evolves through a birth-death scheme. In a given update step, one individual is selected with probability proportional to its fitness to reproduce (encircled with thick solid line), and another individual is chosen for death (encircled with dashed solid line). A reproduction event can result in mutation in one of the daughter cell. In panel A the mutant daughter cell replaces the dead individual, while the parent type daughter cell remains at the parent’s original location. We call this offspring movement. In panel B the parent type daughter cell replaces the dead individual, while the mutant type daughter cell resides at the parent’s original location. We call this parent movement. Although both cases start from the same population, they lead to configurations where the new mutant individual experiences different neighbourhoods.

When parents rather than offsprings move to vacant sites, the same dynamical equation Eq. A1 is recovered for the Bd^*p*^ update rule. The same follows for the dB update rules. Therefore, the deterministic mutation-selection dynamics in a well-mixed population is robust to the choice of update rules, as confirmed also via simulations in Fig. S2 A.

**Fig. S2.**
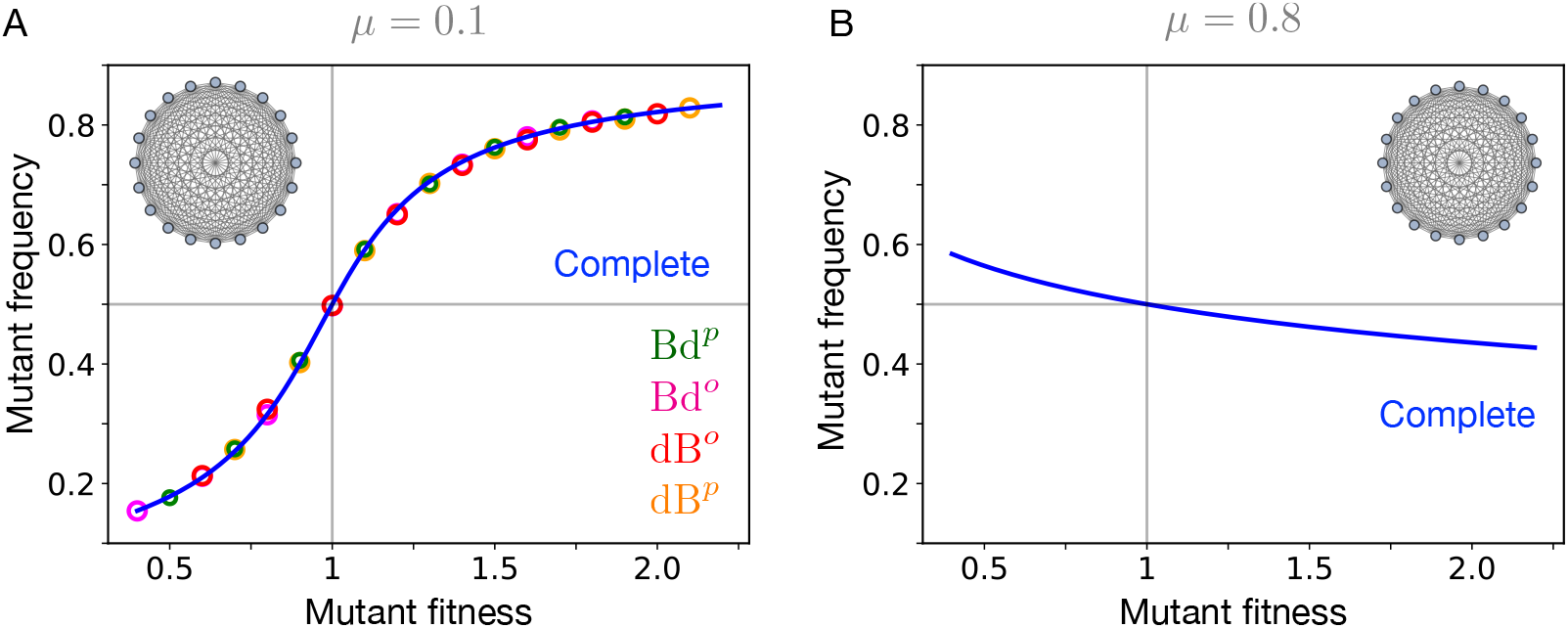
Complete graph. Panel A: Different update rules yield the same mutation-selection balance in a well-mixed population. Here *µ* = 0.1 (*µ <* 1*/*2); circles represent simulation data, while the solid blue line corresponds to the analytical solution from Eq. A2. At low *µ*, selection is a stronger force than mutation, leading to a higher steady-state frequency of beneficial mutants compared to deleterious mutants when competing with the wild-type individuals (*f* = 1). Simulations are performed with *N* = 100. Panel B: For *µ* = 0.8 (*µ >* 1*/*2), mutation dominates over selection, resulting in a higher abundance of deleterious mutants. At *µ* = 0.5 (not shown), the steady-state frequency is equal to 1/2; indicating a perfect balance between natural selection and mutation.

### 2. Star graph

In a star graph, there are two kind of nodes: center and leaf. All *N* −1 leaves are connected to a single center. During an update step, the individual at the center can replace any leaf node, whereas a leaf individual can only replace the center. To study mutation-selection dynamics on the star graph, we track the frequency of mutant-type individuals at the leaves coupled with the dynamics of the central node. The state of the system is defined by *n*_*c*_ ∈ {0, 1}, representing mutant count at the center node and *n*_𝓁_ ∈ { 0, …, *N* − 1}, representing the mutant count at the leaves. We begin with the Bd^*o*^ update scheme.

#### *a*. Bd°

We specify and analyse the transition rates, using the variables *n*_*c*_, *n* ≡ *n*_𝓁_ and 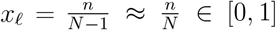. We also introduce the mean fitness for the star graph, which only depends on *x*_𝓁_ and is given by

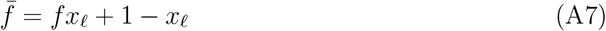

for large *N*. In total, there are four types of transitions that change the state of the central node.

(i) Wild-type center → Mutant center

The center becomes a mutant if:

- A mutant leaf reproduces, and its offspring (without mutation) replaces the center.
- A wild-type leaf reproduces, but its offspring mutates and replaces the center.

Mathematically, the transition probability is

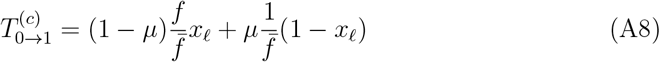

(ii) Mutant center → Wild-type center

The center changes to wild-type if:

- A wild-type leaf reproduces, and its offspring (without mutation) replaces the center.
- A mutant leaf reproduces, but its offspring mutates and replaces the center. The transition probability in that case is

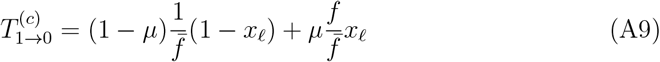

Similarly, the transition probabilities for leaves are

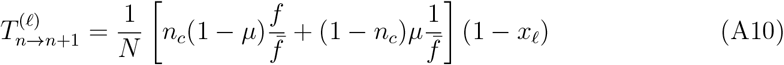

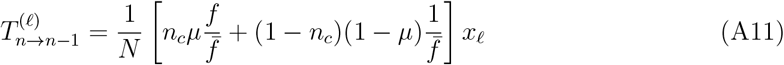

The center’s dynamics is faster than the leaf dynamics by a factor *N*. Thus, we first solve for the stationary center frequency

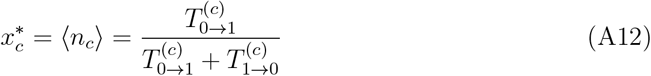

as a function of *x*_𝓁_. Plugging in the transition probabilities (A8,A9), we obtain

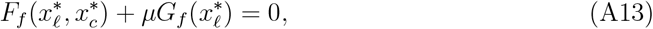

where 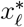 is the stationary mutant leaf frequency and

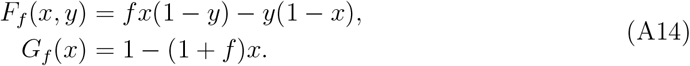

The functions *F*_*f*_ and *G*_*f*_ describe the effect of selection and mutation, respectively, with *f* being the mutant’s fitness.

The expected change in the frequency of mutant-type individuals at the leaves, *x* per update step is given by,

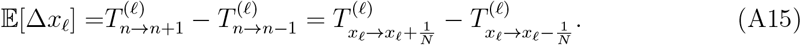

Substituting for the transition probabilities (A10 A11), we obtain

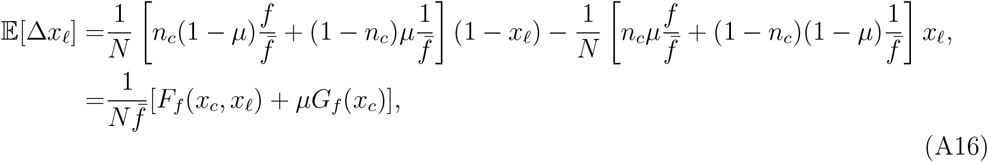

where in the last line, we’ve replaced *n*_*c*_ with its average *x*_*c*_, which is justified by the time-scale separation between the leaf and center dynamics. At steady-state, we have E[Δ*x*_𝓁_] = 0. This together with Eq. A13, leads to following set of equations

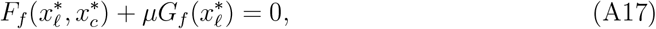

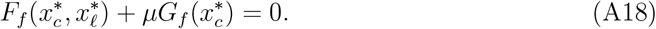

These two equations imply that 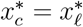.

Moreover, Eq. A17 has the same form as the steady-state equation for the well-mixed population (see Eq. A1). This means that the star graph under Bd^*o*^ update rule exhibits the same mutation-selection balance as the complete graph, with the mutant center frequency equal to the mutant leaf frequency (see Fig. 2A). In other words, 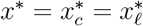. For *µ ≪* 1, we have

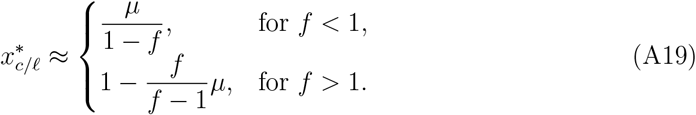

Thus, the star graph under Bd^*o*^ updating has the same mutation load as the well-mixed population. These findings are consistent with the results observed by Yagoobi et. al PRE 2018. However, while Yagoobi et. al [1] assumed 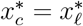 to solve the equations, we recover this equality here through the analysis.

#### *b*. Bd^p^

In this subsection, we focus on the Bd^*p*^ updating where parent individuals replace the dead individuals, while the offspring stays at the position of their respective birth giving individuals. Under this updating scheme, there are four possible transitions that lead to change in the state of the central node.

(i) Wild-type center → Mutant center

The central node, initially occupied by the wild-type, can change to the mutant-type under the following scenarios:

- A mutant leaf node is chosen for reproduction. This is followed by the death of the central node and the parent moving to the center, while the offspring resides at the leaf position.
- The central node is chosen for reproduction, and the offspring mutates. The offspring occupies the center while the parent replaces a random dead leaf individual.

Mathematically, the corresponding transition probability is

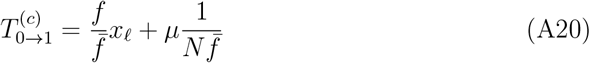

(ii) Mutant center → Wild-type center

Similarly, the central node, initially occupied by the mutant-type, can change to the wild-type under the following scenarios:

- A wild-type leaf node is chosen for reproduction. This is followed by the death of the central node and the parent replacing the dead individual while the offspring stays at the parent’s location.
- The central node is chosen for reproduction with mutation in the offspring, which occupies the center. The parent replaces one of the leaf node individuals.

The transition probability for this case is given by,

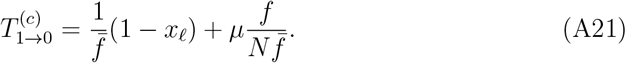

Similarly, the transition probabilities for the leaves are,

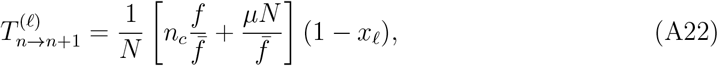

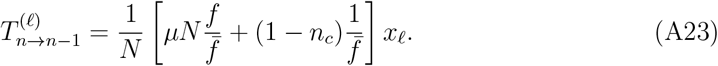

In this updating scheme, the change in the central node’s state is primarily driven by selection, especially for large *N*. The mutation term, has a relatively smaller impact on the central node. On the other hand, the change in the mutant leaf frequency is mostly driven by mutations. Specifically, in Eqs. A22,A23) the product *µN* appears, signalling the strong amplification of mutation in the leaf system. The limit *N* → ∞ can be performed in two ways:

- *N*→ ∞, *µ* → 0 at fixed *U* = *µN* : Then the center dynamics is faster than the leaf dynamics by a factor of *N*, and separation of times scales can be used analogous to the Bd^*o*^ model.
- *N* → ∞ at fixed *µ* ∈ (0, 1): Then the leaf dynamics is dominated by mutation. The limiting rates

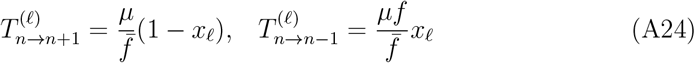

are of the same order as the center rates, but they are *independent of the center state*. As a consequence, we can solve for the leaf frequency and insert the result into (A12) to obtain the center frequency.

Assuming time scale separation, we plug the transition probabilities (A20, A21) into Eq. A12 to obtain an equation satisfied by 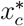 and 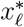

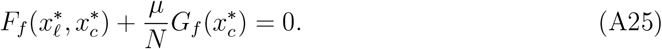

Additionally, the expected change in the mutant leaf frequency 𝔼 [Δ*x*_*𝓁*_] can be derived from the transition probabilities and is given by

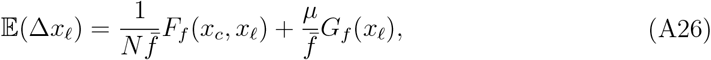

where we have replaced *n*_*c*_ with *x*_*c*_. The term with the prefactor *µ/N* in Eq. A25 can be neglected and thus, in the mutation-selection balance we have,

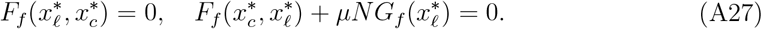

The mutation-selection balance equations can be further simplified into quadratic equations satisfied by 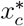 and 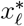

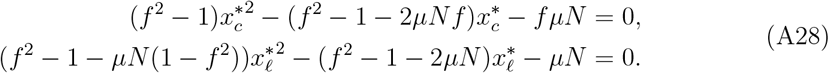

In comparison to the steady-state equations for the complete graph and the star graph under Bd^*o*^ updating, the key difference here is the presence of the term *µN* rather than *µ* in Eq. A27. This makes the effective mutation rate for the leaf nodes equal to *µN*. In this sense, the star graph under Bd^*p*^ updating is an amplifier of mutation.

The solutions to the quadratic equations A28 are

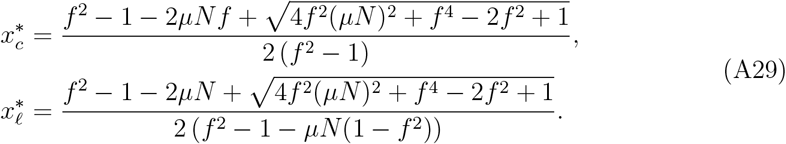

For large *µN*, the solutions can be approximated as

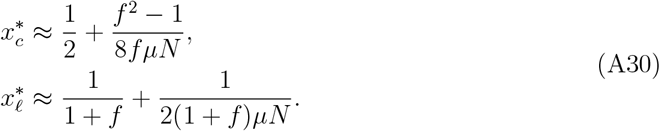

The zeroth order term in the approximation suggests that less fit mutants (lower *f*) will be more abundant in the mutation-selection balance, which is in contrast to a well-mixed population, where fitter mutants dominate. In other words, for the star graph under high mutation rates, the mutation-selection balance profile is qualitatively similar to that of the complete graph when *µ >* 1*/*2. The limit 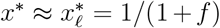 can be derived by considering only the mutation term in Eq. A27 by considering only the mutation term and setting it to zero, 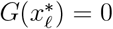. It is equivalent to taking *N*→ ∞ and fixing *µ* to a non-zero value. Thus, the mutational load under Bd^*p*^ updating is

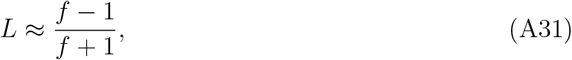

which, for *f ≫* 1, asymptotes to 1 (the upper bound on *L*). Hence, for Bd^*p*^ updating, mutations strongly dominate the evolutionary dynamics over selection.

Now, we ask if there exists a range for *µN* where beneficial mutants are found at higher frequencies than deleterious ones. If such a range exists, there must also be a value of *µN* where 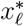 becomes independent of the mutant’s relative fitness. We find that at *µN* = 1, 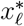 takes a constant value of 1/2 regardless of mutant’s fitness. The series expansion of the mutant frequencies at *µN* = 1 gives

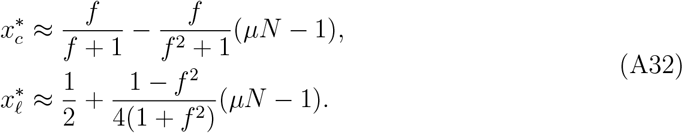

From the above approximation, we see that when *µN <* 1, 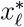 becomes a monotonically increasing function of mutant’s fitness *f*. Conversely, for *µN >* 1, 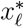 becomes a monotonically decreasing function of fitness.

We conclude this subsection by exploring what happens when both the parent and offspring are allowed to move during the dynamics. In each Bd update step, an offspring moves with probability *ω* while the parent moves with probability 1 −*ω*. A key question is: How much parent movement is needed to strongly amplify mutation? Is it less than 50 percent? From Fig. S3, we find that even a small proportion of parent movement (less than 0.5) is sufficient to generate strong amplification of mutation. This highlights the importance of considering parent movement in evolutionary dynamics.

**Fig. S3.**
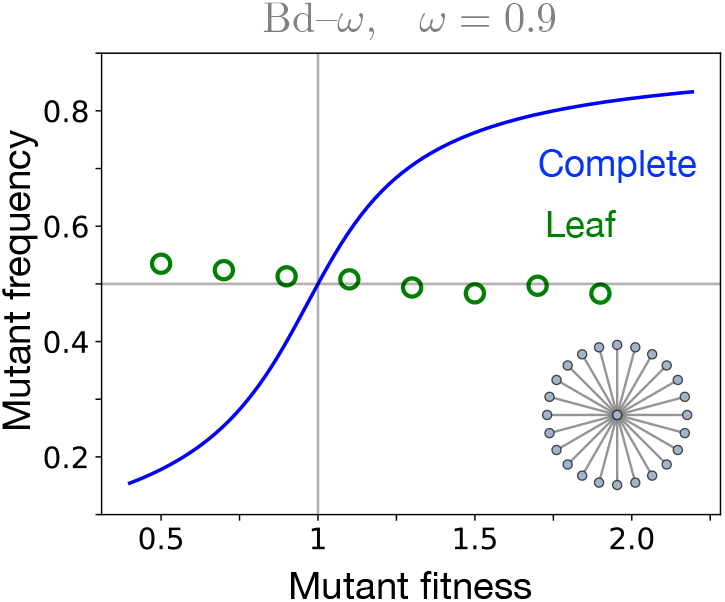
Star graph under Bd–*ω* process. Here, we present simulation results for the Bd–*ω* process, where offsprings move with probability *ω* and parents move with probability 1 − *ω* during reproduction events. With *ω* = 0.9–meaning offspring move 90 percent of the time while parents move 10 percent of the time–we observe a reversal in the mutation-selection profile. This suggests that even a small fraction of parent movement is enough to strongly amplify mutation. Parameters: population size, *N* = 100 with mutation probability *µ* = 0.1.

#### *c*. dB°

We now move to death-Birth update rules, first studying dB^*o*^. Here, death is followed by Birth, with offspring mutating to the other type with probability *µ*. In an update step, an individual is randomly chosen for death with uniform probability, followed by a neighbouring individual (neighbouring dead one) selected with probability proportional to its fitness to reproduce. The offspring replace the dead individual. The transition probabilities for the central node are

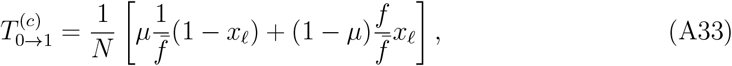

And

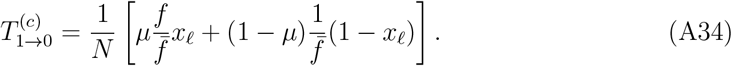

For the leaf nodes, the transition probabilities are

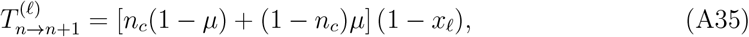

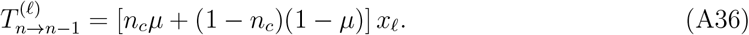

For 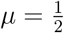, the leaf dynamics decouples from the center, but in general, *n*_*R*_ performs a biased diffusion in a direction determined by *n*_*c*_. The relaxation time of the leaf system is 𝒪 (*N*) and thus of the same order as the relaxation time of the center node according to (A33,A34). Therefore the correlations between *n*_*c*_ and *n*_*R*_ cannot be neglected in this case. To account for these correlations a complete solution for the joint distribution of *n*_*c*_ and *n*_*R*_ would be required, which is not presently available.

Instead, we consider two approximations to obtain the steady-state mutant frequency. In this approach, we neglect any explicit correlations between the central node and leaf nodes. Using Eqs. A33, A34 we substitute for the transition probabilities into Eq. A12 and replacing *n*_*c*_ with *x*_*c*_ gives,

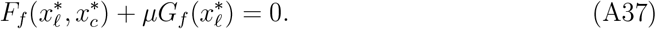

Similarly, the transition probabilities for the leaf nodes (Eqs. B7 and B8) are used to obtain an equation for the expected change in mutant leaf frequency 𝔼 [Δ*x*_*𝓁*_],

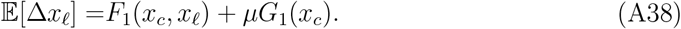

In the steady-state, we have 𝔼 [Δ*x*_*𝓁*_] = 0 which together with Eq. A37 gives

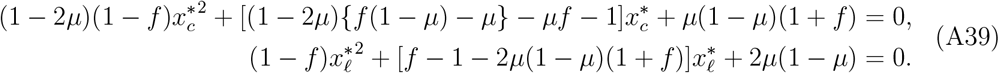

The quadratic equation for 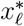 is equivalent to the steady-state equation for the well-mixed population, but with *µ* replaced with 2*µ*(1 −*µ*). In other words, 2*µ*(1 −*µ*) is the effective mutation probability. Thus, under dB^*o*^ updating mutations are amplified over selection.

For *µ* « 1, the approximate solutions to Eq. A39 are given by,

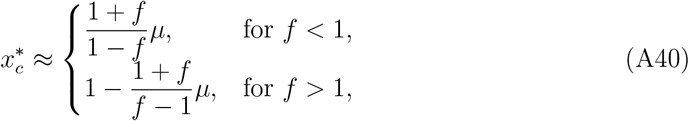

and,

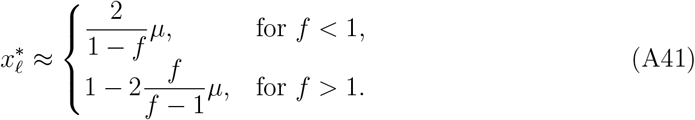

The mutational load *L* is found to be 2*µ*, which is two times the mutation load for the complete graph.

**Fig. S4.**
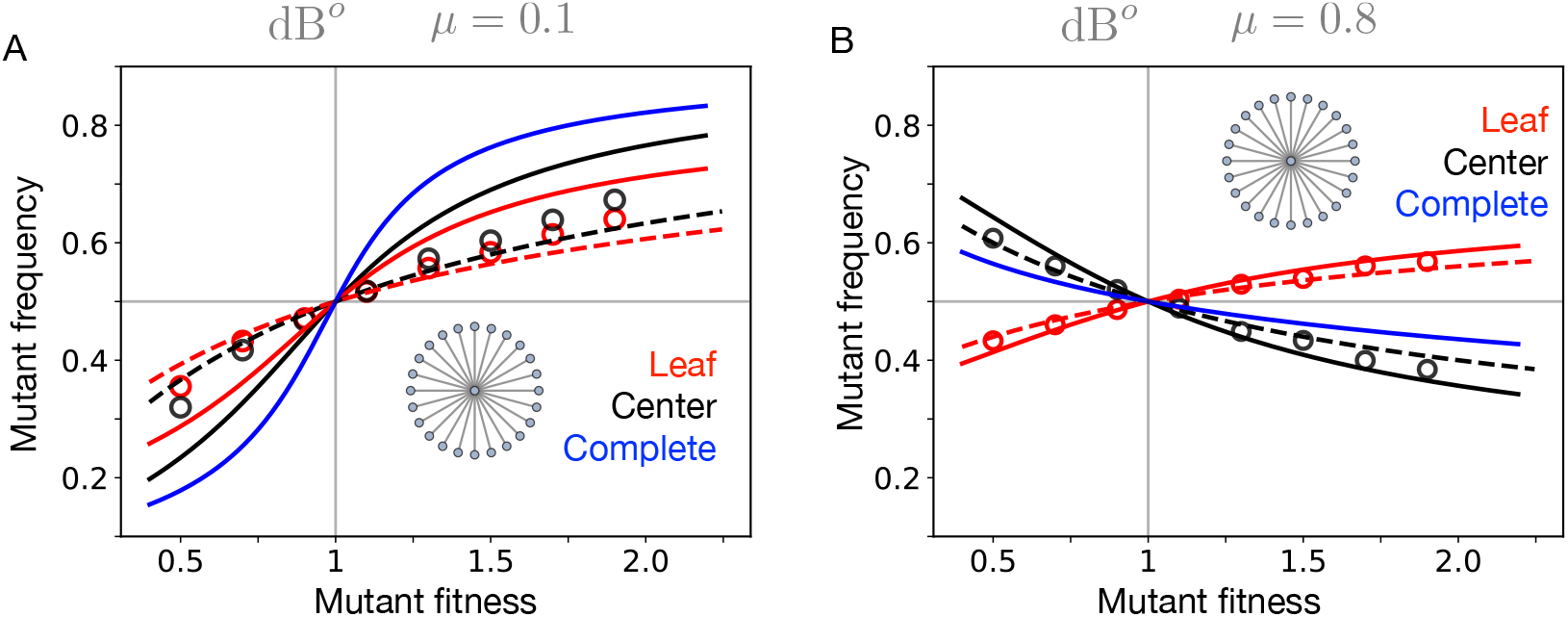
Star graph under dB^*o*^ updating. In panel A, *µ* = 0.1 with *N* = 100 (for simulations). The solid lines represent analytical solution corresponding to Eqs. A39, whereas, the dashed lines correspond to another approximation based on Eqs. A48 and A49. The markers represent simulations. In panel B, *µ* = 0.8. Both approximations, provides bound for the steady-state mean frequencies. The deviations between the approximations and simulations are discussed through Fig. S5. Regardless of the approximation used, the mutation load for the star graph remains higher than the well-mixed population.

Taylor expansion of 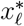 near *µ* = 1*/*2 yields

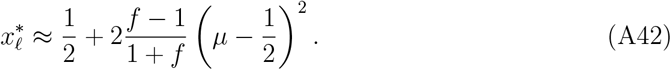

This shows that for *µ* = 1*/*2, the dynamics is neutral, similar to the case for the complete graph. However, the pre-factor of the second term is positive for advantageous mutants and negative for disadvantageous mutants. For higher *µ*, the star graph under dB^*o*^ updating exhibits a greater abundance of beneficial mutants compared to deleterious ones–contrary to the case of complete graph. Furthermore, the mutation-selection balance profiles for *µ* ;S 1*/*2 and *µ* 1*/*2 are identical. Notably, the mutation-selection balance in the leaves for *µ* →0 is the same as for *µ* →1. This symmetry can be seen directly from Eq. A39 where the effective mutation term 2*µ*(1 −*µ*) remains unchanged under the transformation *µ* → 1 −*µ*.

The symmetry in the mutation-selection balance profile can be understood by considering the extreme cases *µ* → 0 and *µ* → 1. Most updates affect the mutant state of the leaves. For *µ* → 0, the center’s state spreads to the leaves. When the central node dies, fitter leaf nodes are more likely to reproduce and replace it, after which the center sends offspring back to the leaves. As a result, higher-fitness individuals dominate the leaf population, as seen in Fig. S4 A and Fig. S5 A.

For *µ* →1, a fitter leaf node is still more likely to replace the center upon its death. However, due to high mutation probability, the new center is typically less fit. The center then sends offspring back to the leaves, reintroducing higher-fitness individuals. This creates a mutation-selection balance profile in the leaves similar to that of the low-mutation case, though opposite for the center. This pattern is evident in Fig. S4 B and Fig. S5 A. Overall, this provides a clear understanding of why the mutation-selection balance profile for the leaves exhibits symmetry at both low and high mutation probabilities.

With that said, the approximation above deviates from the simulations, see Fig. S4. This is because it considers only the marginal frequencies of mutants in the center and leaves, rather than their joint distribution.

We use another approximation that approximately accounts for correlations by assuming a time-scale separation between the center and leaf dynamics. Specifically, we assume that the leaf nodes fully relax to their steady state while the center remains fixed, and vice versa. In this way, the approach effectively assumes maximal correlation between the center and leaves. Under this assumption, the steady-state mutant leaf frequency is given by

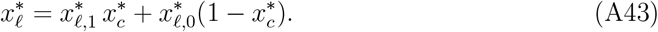

Here, 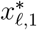 represents the steady-state mutant leaf frequency given that the center is occupied by the mutant type, while 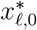 is the steady-state mutant leaf frequency given that the center is occupied by the wild-type. Let us determine 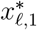. Assuming that the central node is occupied by a mutant-type, the expected change in mutant leaf frequency is given by

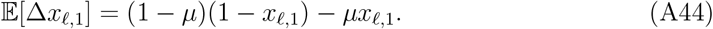

At the steady-state 𝔼 [Δ*x*_*𝓁*,1_] = 0, which gives 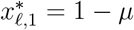, as expected. Similarly, 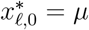. To find 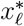, we also need 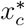. At the steady-state, 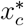 satisfies

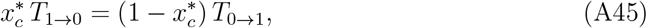

where *T*_1*→*0_ is the probability that the center transitions from mutant-type to wild-type and *T*_0*→*1_ is the probability that the center transitions from wild-type to mutant-type. These transition probabilities are

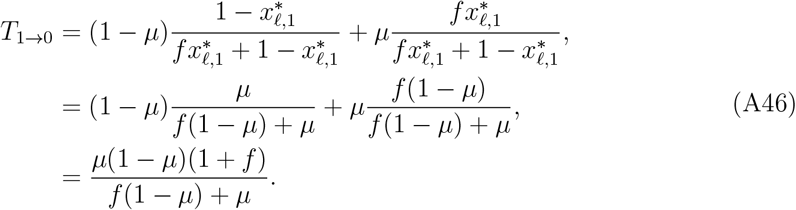

**Fig. S5.**
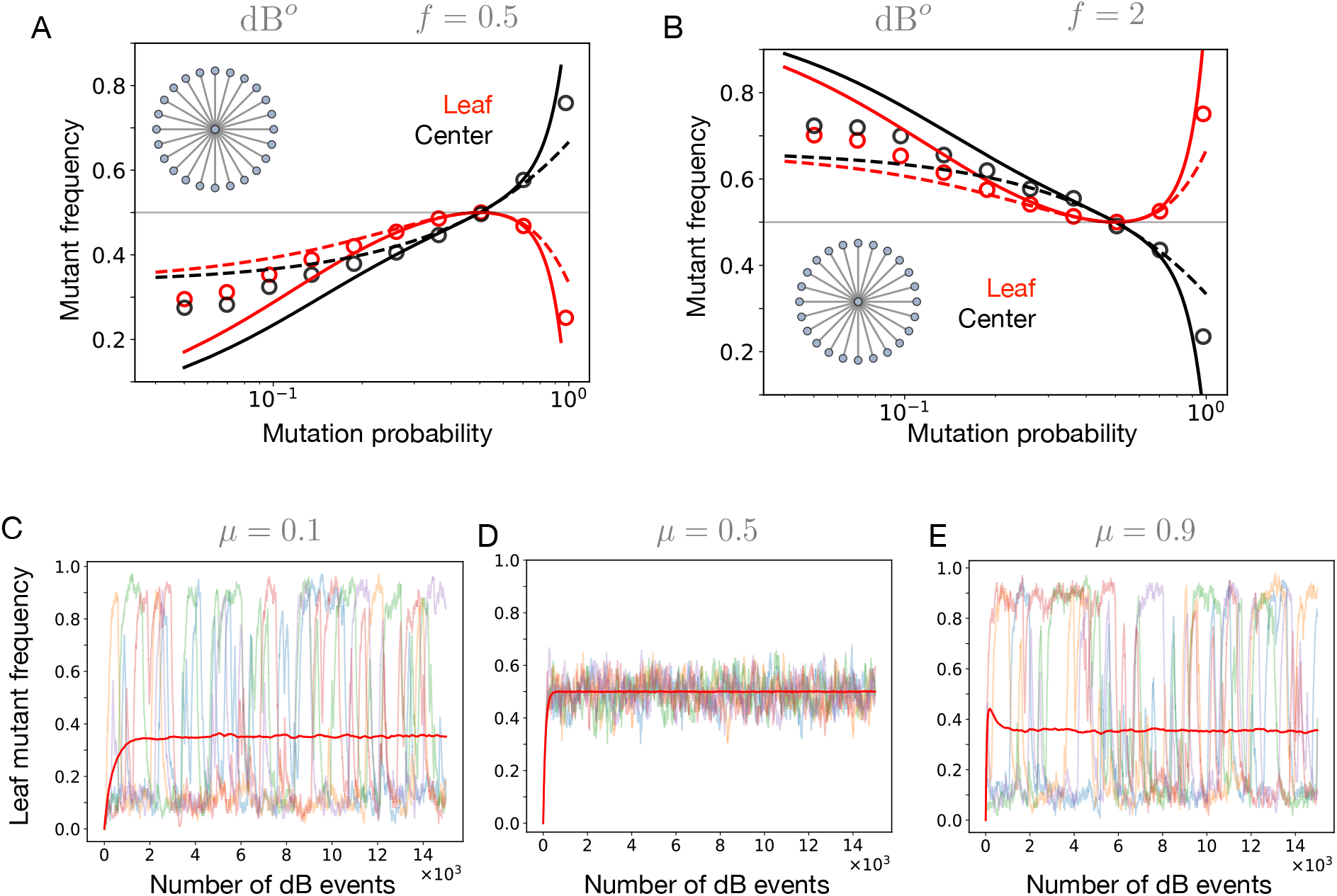
When the mean is not enough: star graph under dB^*o*^ updating. Panel A and B display the deviation between analytical calculations (using two approximations) and simulation results for the steady-state frequencies. The solid lines correspond to the numerical solutions of Eq. A39, while the dashed lines represent the approximations given by Eqs. A48, and A49. Substantial deviations occur at lower and higher *µ* values, but for *µ* near 0.5, there is a good agreement between the calculations and simulations. Additionally, the two different approaches provide bounds for the steady-state frequencies. Panels C, D, and E present the averaged trajectories for leaf mutant frequencies for five independent trajectories. The fluctuations are large for lower and higher *µ* values and do not average out even for large population sizes. In contrast, for *µ* near 0.5, the fluctuations are smaller and symmetric around the mean. Parameters: *N* = 100, with 4000 independent realisations (for panel C,D,E), and the mutant fitness relative to wild-type fitness is 0.5.

Similarly,

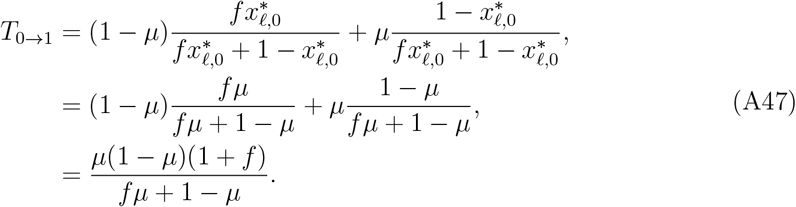

Substituting the transition probabilities into Eq. A45, we find

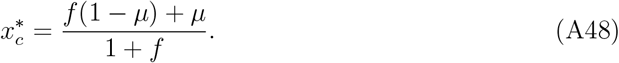

Therefore, using Eq. A43

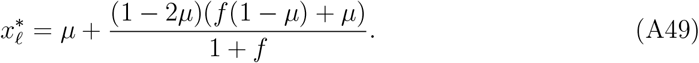

From Fig. S4, we find that these two approximations provide upper and lower bounds to the steady-state average frequencies.

#### *d*. dB^p^

For the dB^*p*^ update, each step consists of a death event followed by a birth event. However, unlike the previous case, the parent moves to the vacant site instead of offspring replacing the dead individual. The transition probabilities for the center node are,

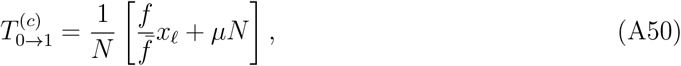

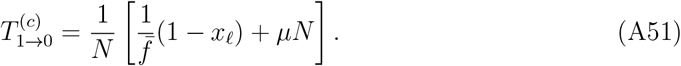

Similarly, the transition probabilities for the leaf nodes are,

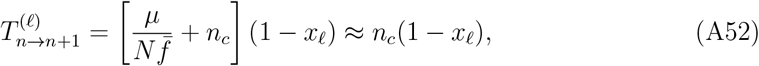

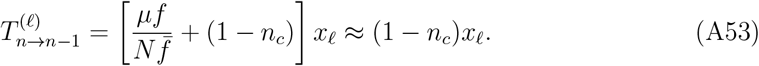

In Eqs. A50,A51 the product *µN* appears, signalling the strong amplification of mutation whereas the leaf dynamics is primarily driven by natural selection.

The limit *N* → ∞ in (A50,A51) can be performed in two ways:

- *N* → ∞, *µ* → 0 at fixed *U* = *µN* : Then the relaxation time for the center dynamics is of the same order as the leaf dynamics. As a result, the center and leaf dynamics become correlated, similar to the case of dB^*o*^ dynamics.
- *N*→ ∞ at fixed *µ* ∈ (0, 1): Then the center dynamics is dominated by mutation and the center relaxes 𝒪(*N*) times faster than the leaves. Consequently, the time-scale separation approximation used in Bd update schemes is expected to hold well in this regime.

Assuming *µ* is fixed and taking large *N* limit, the resulting time-scale separation allows us to compute the steady state of the center. Using Eqs. A50,A51, A12, we find the 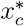 satisfies the equation

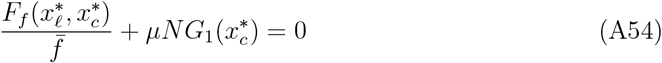

Similarly, from Eqs. A52,A53 we obtain 𝔼 [Δ*x*_*𝓁*_] = *F*_1_(*x*_*𝓁*_, *x*_*c*_). In the steady-state yields 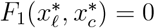, implying that 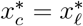. Using this equality in Eq. A54, we find

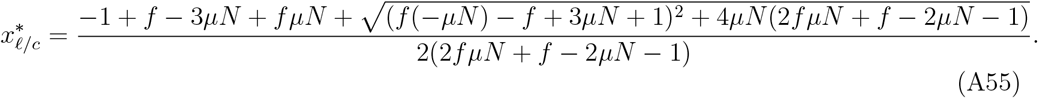

Approximating the exact solution for large *µN*, we obtain

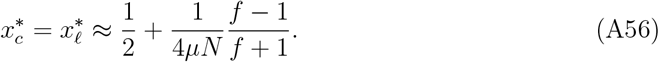

For *µN ≫*1, we the zeroth order term dominates, yielding 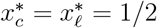. This leads to,

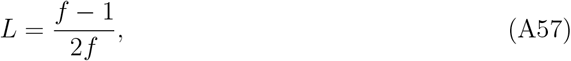

which further asymptotes to 1*/*2 for *f* 1. Under dB^*p*^ updating, the dynamics on the star graph become effectively neutral, see Fig. S6 A. The effects of selection are eliminated by mutations, making the mutation-selection balance primarily driven by mutations. To build intuition for this result, consider the update process: in most steps, a leaf node is selected for death, followed by reproduction from the central node. Since, during mutation, the offspring remains at the birth-giving node, the central node’s probability of being occupied by a mutant stabilizes at 1*/*2, assuming no mutation bias. Because the center determines the fate of the leaf nodes, their mutant frequency mirrors that of the center. Consequently, the steady-state mutant frequency in the leaves also settles at 1*/*2.

As *µN* decreases, the simulations begin to deviate from our approximation, as shown in Fig. S6 B C. This deviation arises because the time-scale separation assumption becomes less accurate due to increased correlations in the central and leaf dynamics.

**Fig. S6.**
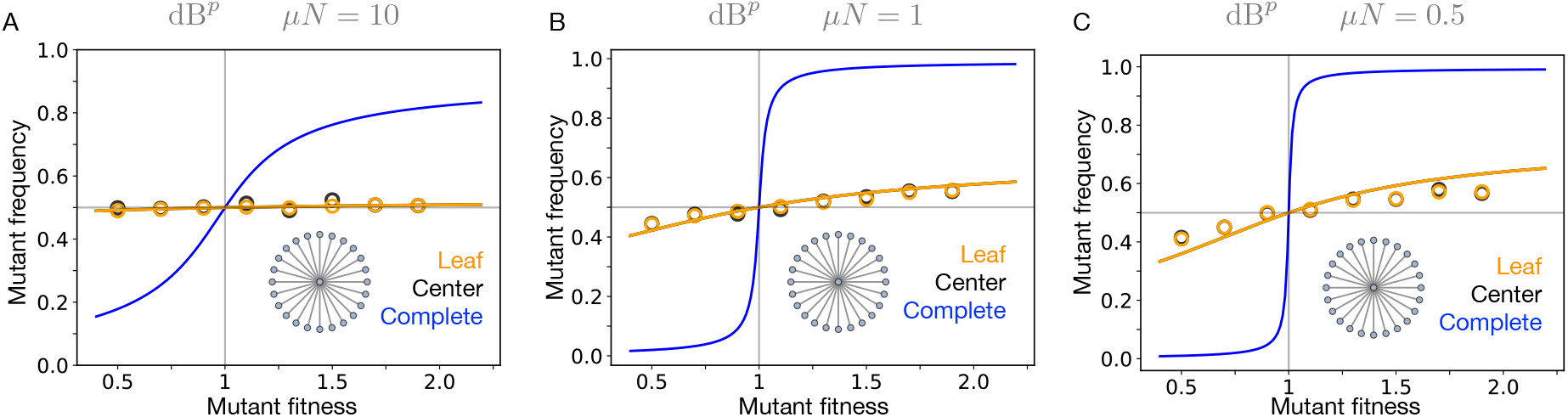
Star graph under dB^*p*^ updating. In Panel A (*µN* = 10, with *µ* = 0.1 and *N* = 100), the mutation-selection balance on the star graph aligns with what is expected for neutral evolution. The steady-state mutant frequency remains at 1*/*2, independent of the mutant’s fitness. In Panel B (*µN* = 1, with *µ* = 0.01 and *N* = 100), we observe a departure from the neutral steady-state limit. As *µN* decreases, selection begins to take effect, causing deviations from the expected 1*/*2 frequency. In Panel C (*µN* = 0.5, with *µ* = 0.005 and *N* = 100), simulation results further deviate from the analytical approximation. This deviation arises due to the correlations between the center and leaf nodes, which become more pronounced at lower mutation probabilities. As *µ* decreases, the assumption of independence between the center and leaf dynamics breaks down, leading to noticeable differences between simulations and analytical predictions.

### 3. Barabási–Albert graph

Here, we present simulation results for the Bd^*p*^ updating on Barabasi–Albert (BA) graphs generated using the non-linear preferential attachment (NLPA) model [2–4]. The graph construction begins with *m*_0_ initial nodes. At each time step, a new node is added and connected to *m* existing nodes (with *m* ≤*m*_0_). These *m* nodes are sampled with a probability proportional to 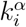, where *k*_*i*_ is the degree of node *i* and *α* is a positive constant. This process continues until the graph contains *N* nodes.

- For *α* = 1, the NLPA model recovers the standard Barabasi–Albert graph with a scale-free degree distribution.
- For *α <* 1, the resulting graph exhibits a stretched exponential degree distribution.
- For *α >* 1, the attachment process leads to a highly centralized star-like structure in which most nodes connect to a single hub.

**Fig. S7.**
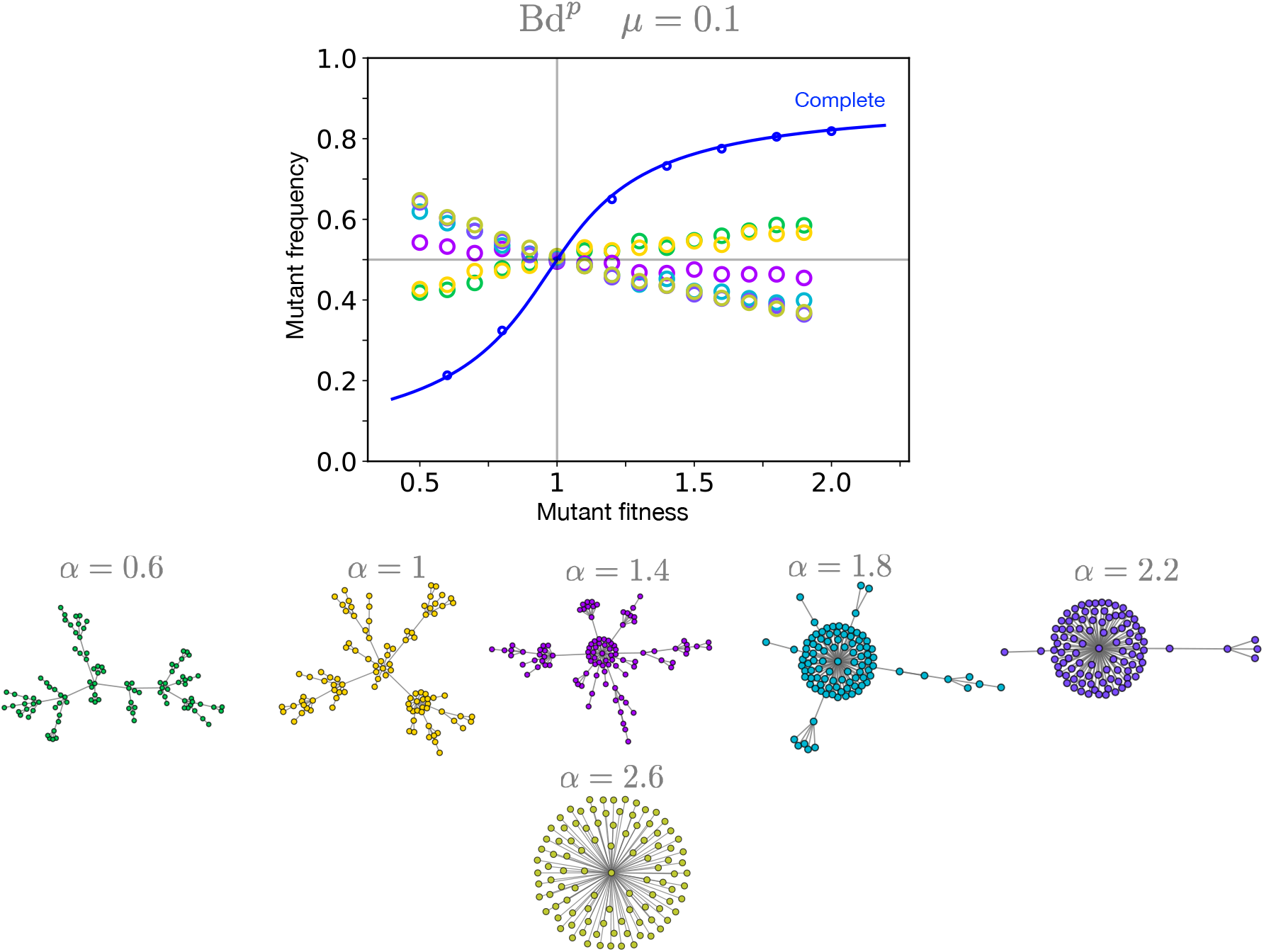
Non-linear preferential attachment model with Bd^*p*^ updating. Simulation results for the NLPA model under Bd^*p*^ updating are shown for various values of *α*. For *α* = 1, the model generates the standard Barabási–Albert (BA) graph. When *α <* 1, the resulting graphs exhibit stretched exponential degree distributions, while *α >* 1 leads to star-like structures. Among the simulated *α* cases, we observe a reversal in the mutation-selection profile for *α >* 1 where deleterious mutants tend to be in a higher frequencies than the beneficial mutants. Parameters: *N* = 100, *µ* = 0.1, and *m* = 1 (number of nodes to which new nodes connect in the graph generation algorithm).

## Appendix B: Mode of mutation: spontaneous

In this section, we analyze update schemes where mutations occur spontaneously in the population, rather than being coupled to birth events. As usual, we start with the reference case, the complete graph.

### 1. Complete graph

In each update step, mutation occurs with probability *µ*. If mutation happens, a randomly chosen individual changes its state. If no mutation occurs (with probability 1 − *µ*), a birth-death update takes place. The expected change in mutant frequency is then given by

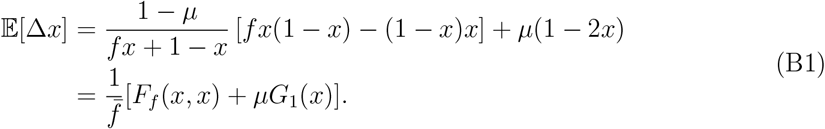

Setting 𝔼 [Δ*x*] = 0, the exact solution for the mutation-selection balance is,

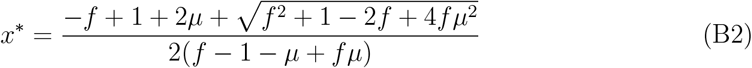

For small *µ, x*^*\**^ approximates to

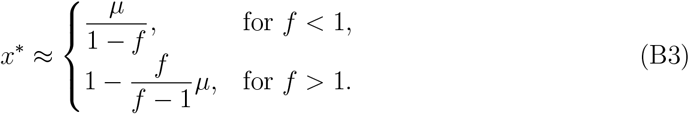

This low *µ* approximation for the complete graph with spontaneous mutations yields the same mutation-selection balance as the case of mutation coupled to birth, see Eq. A3 and Fig. S8 A. Consequently, the mutational load *L* is also the same. However, unlike the case with mutation coupled to birth, there is no reversal in the mutation-selection balance profile here, see Fig. S8 B. Specifically, as the mutation probability increases, the mutation-selection balance flattens out, with *x*^*\**^ = 1*/*2 at *µ* = 1. Near *µ* = 1, Eq. B2 can be approximated as

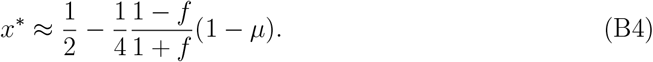

The zeroth order can be obtained directly from Eq. B1 by setting *G*_*f*_ (*x*) = 0. At *µ* = 1, selection is absent and mutation is the only active evolutionary force. Since, there is no explicit mutation bias, the steady-state mutant frequency at *µ* = 1 is 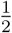. As before, the complete graph is robust to the choice of update rule - Bd or dB. With this established, we now proceed to analyze the star graph.

### 2. Star graph

#### *a*. Bd

The transition probabilities for the central node are,

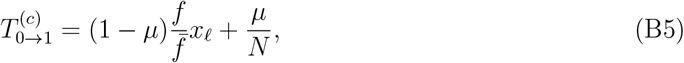

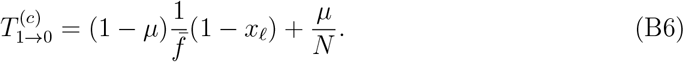

The transition probabilities for the leaves are,

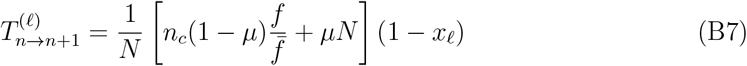

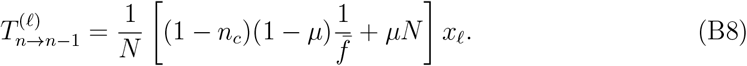

Similar to Bd^*p*^, the product *µN* appears again in the transition probabilities for leaf nodes. We can proceed in the same way as we did in case for Bd^*p*^. Substituting Eqs. B5 and B6 into Eq. A12, we obtain an equation satisfied by 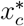

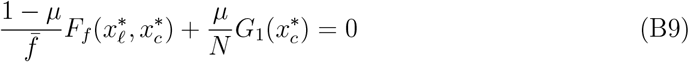

Similarly, the equation for the expected change in mutant leaf frequency 𝔼 [Δ*x*_*𝓁*_] is

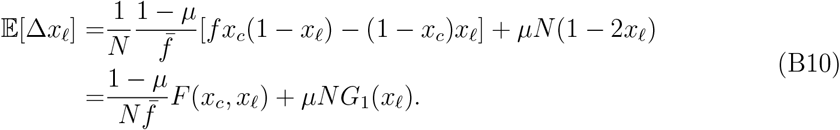

In the steady-state, 𝔼 [Δ*x*_*𝓁*_] = 0, which gives

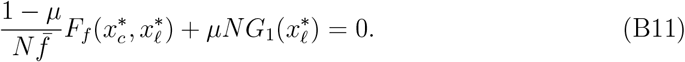

As before, we neglect the term with a pre-factor *µ/N* in Eq. B9 and the term with a pre-factor 1*/N* in Eq. B11. This simplification gives us 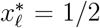 and 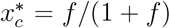. Consequently, the mutation load is 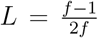. The steady-state mutant leaf frequency, 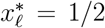, is the same as what we would expect in a neutral dynamics scenario. In other words, mutations eliminate the effect of selection on the star graph. This is analogous to the result found for the well-mixed population at *µ* = 1. Thus, the star graph under Bd updating with spontaneous mutation behaves as an amplifier of mutation, as seen in Fig. S9. The intuition behind this result is as follows: The mutant leaf frequency changes either when the center is chosen for reproduction or when a mutation occurs at the leaf nodes. For large population sizes, the frequency change is primarily driven by mutations. As a result, we can effectively ignore selection, with mutations completely dictating the dynamics. Since there is no explicit mutation bias, the steady-state mutant leaf frequency becomes1*/*2.

The star graph amplifies selection in the weak mutation regime. However, at higher mutation rates, this amplification is lost, and mutation dominates. Overall, whether mutations are coupled to birth or not, the star graph amplifies mutation under Bd updating.

#### *b*. dB

For dB updating with spontaneous mutations, the transition probabilities for the central node are,

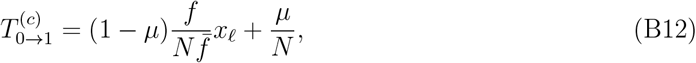

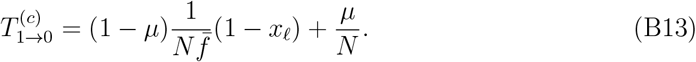

The transition probabilities for the leaves are,

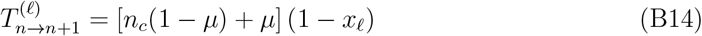

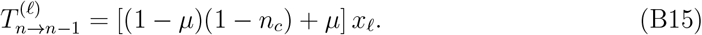

The relaxation time of the leaf system is 𝒪(*N*) and thus of the same order as the relaxation time of the center node according to (B12,B13). Thus, the correlations between *n*_*c*_ and *n*_.*e*_ cannot be neglected.

However, to estimate the mutation load, we proceed by ignoring these correlations. Assuming time-scale separation between the center and leaf node dynamics, the steady-state average mutant occupancy probability for the central node satisfies

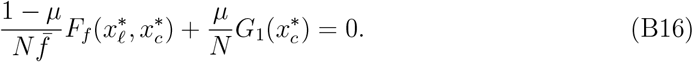

Similarly, the equation for the expected change in mutant leaf frequency 𝔼 [Δ*x*_*𝓁*_] is

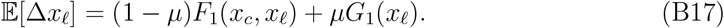

Setting 𝔼 [Δ*x*_*𝓁*_] = 0, and approximating 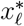 for low *µ*, we obtain

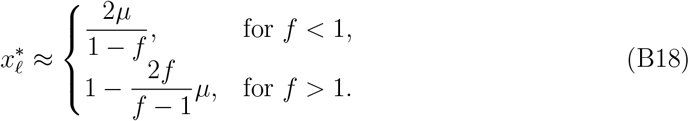

By comparing with Eq. B3, we conclude that the star graph under dB updating with spontaneous mutations acts an amplifier of mutation, with an effective mutation probability of 2*µ*. The mutational load is approximately 2*µ*. The analytical solutions deviate from the simulations at low *µ* values, see Fig. S10 A. This deviation is again due to strong fluctuations in the leaf mutant frequency, which average out at higher mutation probabilities, as seen in Fig. S10 B and C.

**Fig. S8.**
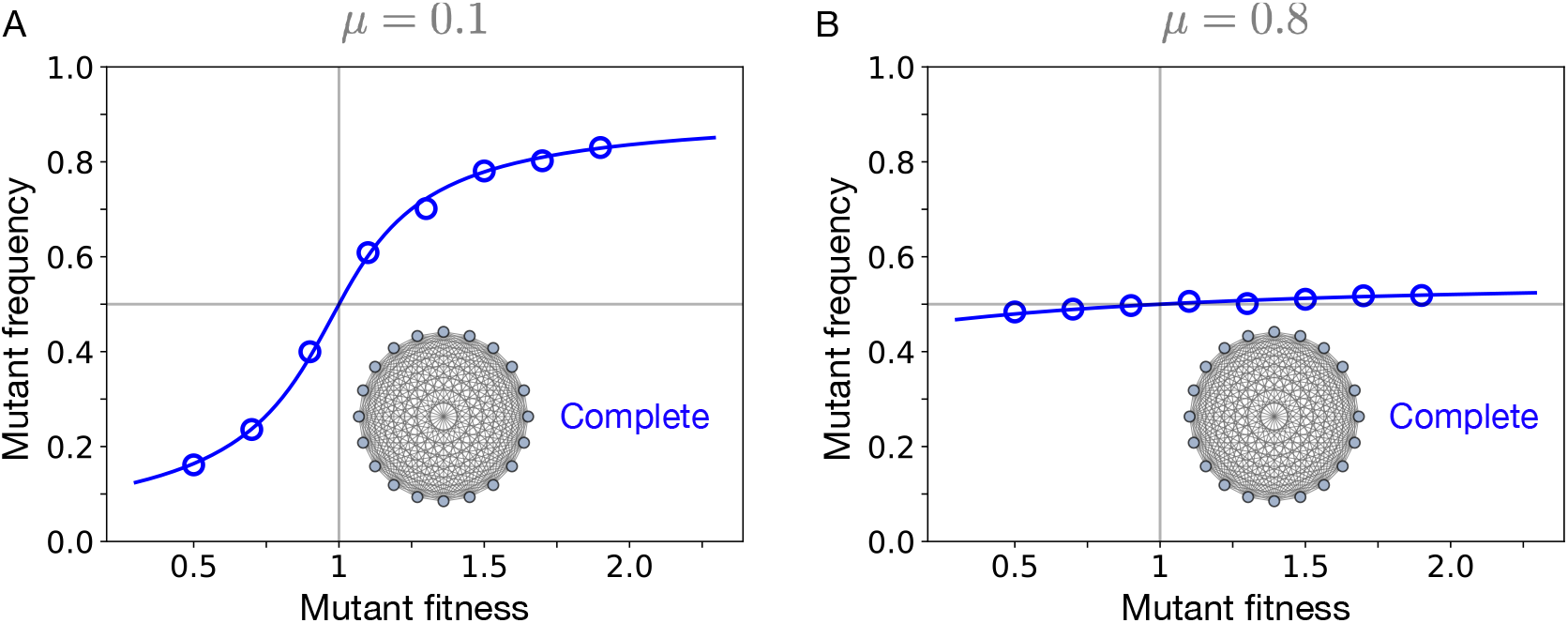
Spontaneous mutations and the complete graph. The average steady-state mutant frequencies as a function of mutant relative fitness are shown, with the wild-type fitness set to one. Mutations occur spontaneously with probability*µ*. In panel A, *µ* = 0.1, while in panel B, *µ* = 0.8. Solid lines represent the exact solution (Eq. B2) and symbols corresponds to simulation results. For low mutation probabilities, the mutant frequency profile closely resembles that of mutation coupled to birth (Fig. S2). However, for higher mutation probabilities, there is no reversal in the mutant frequency profile.

**Fig. S9.**
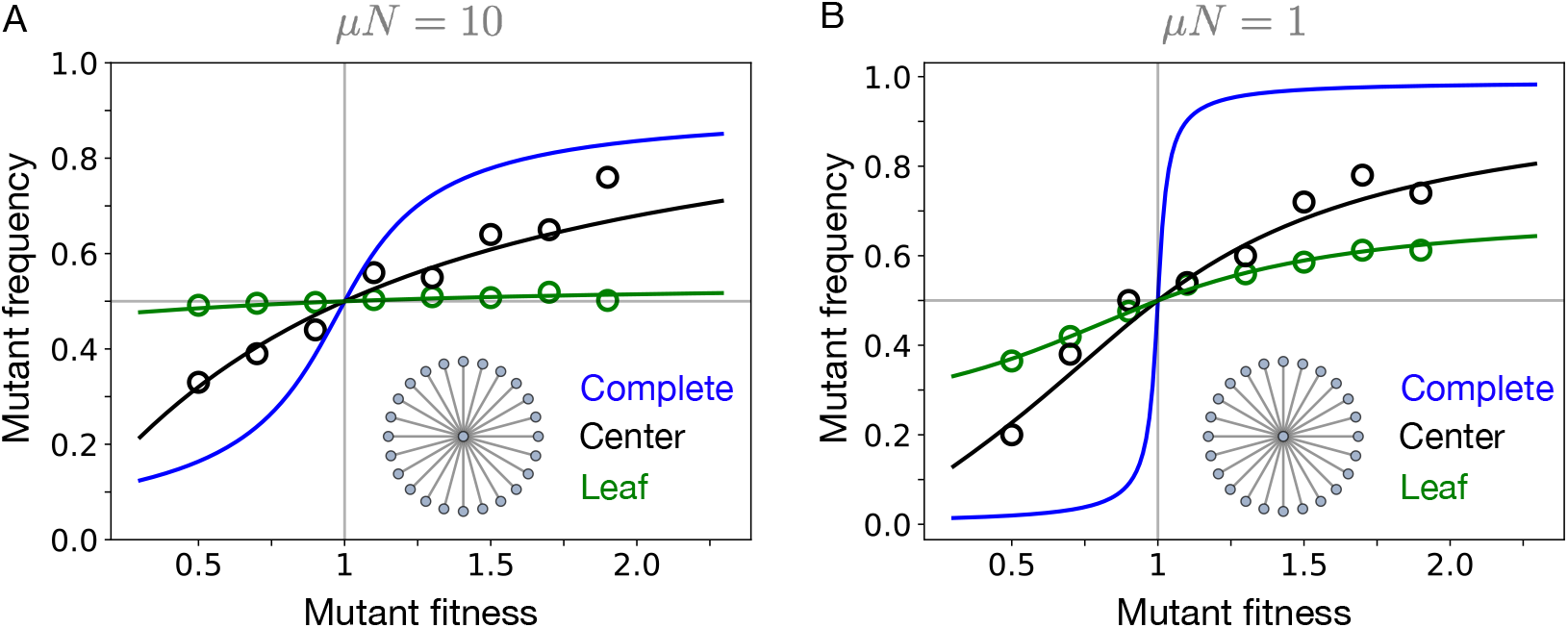
Spontaneous mutations and the star graph under Bd updating. The average steady-state mutant frequencies as a function of mutant relative fitness are shown, with the wild-type fitness set to one. In each update step, a mutation occurs with probability *µ*, or a Bd step takes place with probability 1 − *µ* on the star graph. Solid lines represent numerical solutions of the coupled system (Eq. B11), while symbols correspond to simulation results. In panel A, the mutation probability is *µ* = 0.1 with *N* = 100, whereas in panel B, *µ* = 0.01 making *µN* = 1.

**Fig. S10.**
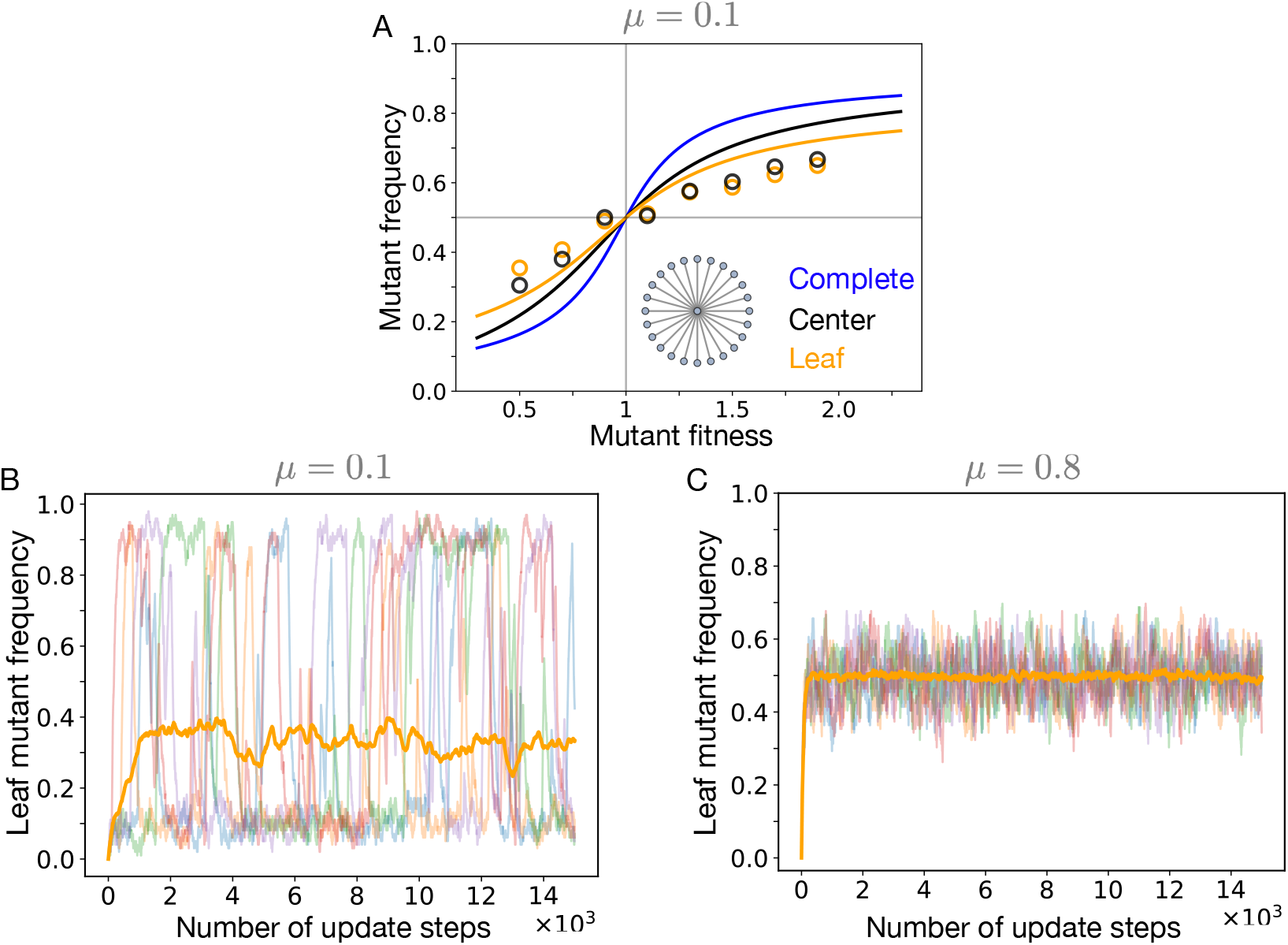
Spontaneous mutations and the star graph with dB updating. In panel A, the steady-state frequencies are shown for the star under dB updating with mutations appearing spontaneously. The mutation probability *µ* = 0.1. Solid lines correspond to numerical solution of the coupled system of Eqs. B16 B17, and symbols represent simulations. While both numerical and simulation suggests that the star graph has higher mutational load than the well-mixed population, the analytical solutions deviate from the simulations. This is because for lower *µ* values the fluctuations in the leaf mutant frequency are strong as shown in panel B, with orange curve representing the mean mutant leaf frequency averaged over 100 independent realisations and background shows five such independent leaf mutant trajectories. As shown in panel C, for higher *µ* (here *µ* = 0.8), the fluctuations are small and averages out. Parameters: (panel A, B and C) *N* = 100, for panel B and C, the mutant fitness relative to wild-type fitness is 0.5.

